# A target enrichment probe set for resolving the flagellate plant tree of life

**DOI:** 10.1101/2020.05.29.124081

**Authors:** Jesse W. Breinholt, Sarah B. Carey, George P. Tiley, E. Christine Davis, Lorena Endara, Stuart F. McDaniel, Leandro G. Neves, Emily B. Sessa, Matt von Konrat, Sahut Chantanaorrapint, Susan Fawcett, Stefanie M. Ickert-Bond, Paulo H. Labiak, Juan Larraín, Marcus Lehnert, Lily R. Lewis, Nathalie S. Nagalingum, Nikisha Patel, Stefan A. Rensing, Weston Testo, Alejandra Vasco, Juan Carlos Villarreal, Evelyn Webb Williams, J. Gordon Burleigh

## Abstract

**Premise of the study:** New sequencing technologies enable the possibility of generating large-scale molecular datasets for constructing the plant tree of life. We describe a new probe set for target enrichment sequencing to generate nuclear sequence data to build phylogenetic trees with any flagellate plants, comprising hornworts, liverworts, mosses, lycophytes, ferns, and gymnosperms.

**Methods and Results:** We leveraged existing transcriptome and genome sequence data to design a set of 56,989 probes for target enrichment sequencing of 451 nuclear exons and non-coding flanking regions across flagellate plant lineages. We describe the performance of target enrichment using the probe set across flagellate plants and demonstrate the potential of the data to resolve relationships among both ancient and closely related taxa.

**Conclusions:** A target enrichment approach using the new probe set provides a relatively low-cost solution to obtain large-scale nuclear sequence data for inferring phylogenetic relationships across flagellate plants.

## INTRODUCTION

For the first ~300 million years following plants’ movement and adaptation to land, Earth’s terrestrial flora consisted of flagellate plants, or plants with mobile flagellate male gametes (i.e., spermatozoids). The modern descendants of these lineages that have retained flagellate sperm include the hornworts, liverworts, mosses, lycophytes, ferns, and some gymnosperms, which comprise approximately 30,000 extant species. During the evolution of these groups, numerous anatomical innovations arose, including stomata, vascular tissue, roots and leaves, lignified stems with secondary growth, and seeds. Collectively, these plants hold the keys to understanding the early evolution of these and other critical features of modern land plant diversity, which is overwhelmingly represented by non-flagellate angiosperms. Despite their long evolutionary history, the phylogenetic relationships among many flagellate plant taxa remain poorly understood, and the lack of a consistent molecular toolkit makes resolving these relationships difficult.

Analyses of large numbers of nuclear loci can provide the power to resolve difficult phylogenetic relationships and the ability to address patterns of lineage sorting and reticulate evolution. Recent analyses of single-copy nuclear genes from transcriptome data have provided insights into backbone relationships among flagellate plants (Wickett et al., 2014; Shen et al., 2018; Qi et al., 2018; One Thousand Plant Transcriptomes Initiative, 2019). However, transcriptome sequencing requires access to freshly collected tissue and often is expensive and impractical, and many loci are either not useful for phylogenetics or only expressed in specific tissues or stages of development. Therefore, transcriptomic sequencing approaches may not be feasible for building large-scale phylogenetic trees (see McKain et al., 2018). Target enrichment methods use short RNA probes, corresponding to selected loci, to bind to DNA from sequencing libraries. The bound DNA is then sequenced, while much of the unbound DNA is discarded (Gnirke et al., 2009; Cronn et al., 2012; Weitemier et al., 2014). Target enrichment approaches can be used to obtain data from hundreds of phylogenetically informative nuclear loci at relatively low cost. These approaches also appear to work well with low-quantity, potentially degraded DNA samples, like those extracted from herbarium specimens (see Brewer et al., 2019, Forrest et al., 2019). Target enrichment approaches have been used to generate nuclear datasets to resolve relationships within several flagellate plant clades, including mosses (Liu et al., 2019; Medina et al., 2019), ferns (Wolf et al., 2018), and pines (Gernandt et al., 2018; Montes et al., 2019). However, generating large nuclear datasets for phylogenetic inference among most flagellate plant taxa remains challenging.

In this study, we leveraged recent transcriptome and whole genome sequence data to design a “universal” probe set that enables target enrichment sequencing across all flagellate plant lineages. The probes were designed to target 451 relatively conserved exons in single or low copy nuclear loci. Furthermore, the target enrichment protocol typically also yields sequence data from the more variable flanking regions that may be useful to resolve relationships among closely related taxa. We demonstrate the target enrichment protocol using representative from all major flagellate plant lineages and provide an analytical pipeline to process the resulting data.

## METHODS

### Probe design

We designed target enrichment probes to cover all flagellate plant groups, including mosses, liverworts, hornworts, lycophytes, ferns, and gymnosperms, using existing genomic and transcriptomic data. We designed probes to cover conserved exons in single (or low) copy nuclear loci identified by the 1KP initiative (DOI 10.25739/8m7t-4e85; Carpenter et al., 2019), which assembled transcriptomes from 1,173 green plant species. We examined available genome sequences from land plant taxa in the 1KP alignments to identify exons that were at least 120 base pairs (bp) in length that belong to the single copy loci identified by 1KP. We used a pairwise BLAST of selected exons to find those shared across multiple genomes that were at least 120 bp long and had at least 65% average pairwise identity. Only regions represented across multiple genomes, suggesting conservation of exon content across land plants, were used to design probes. For the probe kit, we identified the best 451 loci (i.e., exons) that have splice sites conserved across multiple genomes. In some cases, multiple exons used in the probe set are found within the same gene; in total, the 451 exons we used are found in 248 genes (see Supplemental Table 1). We aligned and then cut these loci out of the 1KP alignments that included only the flagellate plant taxa. We clustered the cut sequences for each locus at 90% similarity and took the centroid sequence of each cluster. We designed the probe set from these sequences with a 2x tiling density. The resulting *GoFlag 451* probe set consists of 56,989 probes covering 451 loci and is available on Dryad (https://doi.org/10.5061/dryad.7pvmcvdqg). The term GoFlag refers to the Genealogy of Flagellate plants project, which was funded through the NSF Genealogy of Life (GoLife) program.

To test whether the 451 exons would be phylogenetically informative across land plants, we extracted these exons from the 1KP translated nucleotide alignments and removed sequences from non-land plants from the alignments. We concatenated the exon alignments into a supermatrix and ran a maximum likelihood (ML) search with 100 nonparametric bootstrap (BS) replicates using RAxML 8.2.10 with the GTR CAT model (Stamatakis, 2014). Alignments for this analysis also are available on Dryad (https://doi.org/10.5061/dryad.7pvmcvdqg).

### Taxon Selection

We assembled a collection of 188 samples for our pilot study (Supplemental Table 2). These include representatives of major clades within hornworts (14), liverworts (46), mosses (48), lycophytes (16), ferns (48), and gymnosperms (16). Within these groups we also included some sets of closely related taxa (e.g., congeners) to test the probe set’s ability to resolve close relationships (see Supplemental Table 2 for voucher information). Some of these samples came from herbarium specimens, while others were from recently collected silica dried tissue. We extracted DNA using a cetyl trimethyl ammonium bromide (CTAB) extraction, described in Doyle and Doyle (1987), modified for 2-mL extractions, using a Genogrinder 2010 mill (SPEX CertiPrep, Metuchen, NJ), and with 2.5% polyvinylpyrrolidone and 0.4% beta-mercaptoethanol, and two rounds of chloroform washes followed by an isopropanol precipitation and an ethanol wash. To remove RNA contamination, between chloroform washes we added 0.2uL of RNase A (Qiagen, Valencia, CA, USA) to each sample.

### Sequence Capture and Sequencing

The library construction, target enrichment, and sequencing were done by RAPiD Genomics (Gainesville, FL, USA). After a bead-based DNA cleanup step, DNA was normalized to 250 nanograms (ng) and mechanically sheared to an average size of 300 base pairs (bp). We constructed next-generation libraries by repairing the ends of the sheared fragments followed by the addition of an adenine residue to the 3’-end of the blunt-end fragments. Next, we ligated barcoded adapters suited for the Illumina sequencing platform to the libraries. Ligated fragments were PCR-amplified using standard cycling protocols (e.g., Mamanova et al. 2010). We pooled 16 barcoded libraries equimolarly to a total of 500 ng for hybridization. Target enrichment was performed using the custom designed probes and protocols as suggested by Agilent (Palo Alto, California, USA). After enrichment, samples were re-amplified for additional 6-12 cycles. All enriched samples were sequenced using an Illumina HiSeq 3000 with paired-end 100 bp reads. The sequence reads were deposited in the NCBI sequence read archive (SRA; see Bioproject PRJNA630729).

### Bioinformatic and phylogenetic analyses

Targeted nuclear exon loci were recovered from enriched Illumina data using a modified version of the iterative baited assembly pipeline described by Breinholt et al. (2018). Our six-step pipeline, with all scripts and necessary input files, is available in Dryad (https://doi.org/10.5061/dryad.7pvmcvdqg). In step 1 (*trim reads*), adapters and bases with Phred scores less than 20 were trimmed from paired-end reads with Trim Galore! version 0.4.4 (https://www.bioinformatics.babraham.ac.uk/projects/trim_galore/). Only pairs of reads in which both the forward and reverse read were at least 30 bp long were retained for assembly. In step 2 (*assembly*), the targeted loci were assembled using iterative baited assembly (IBA) implemented in a previously published Python script (IBA.py; http://datadryad.org/resource/doi:10.5061/dryad.rf7g5.2; Breinholt et al., 2018). For each locus, the script first finds raw reads with significant homology to the probe region based on the reference transcriptome sequences from OneKP data and whole genome sequences (Supplementary Material) using USEARCH version 7.0 (Edgar, 2010) and then performs an iterative *de novo* assembly with the subset of reads for each locus with BRIDGER version 2014-12-01 (Chang et al., 2015). In the IBA script, we set the BRIDGER kmer size parameter to 25 and the minimum depth of coverage for the kmers to be included in the assembly to 10. We set the number of IBA iterations to 3 in order to extend the assembly beyond the probe regions. In step 3 (*probe trimming*), we separate the probe region sequences to be used in the next step to assess orthology and format the output.

Although the probes were designed from exons in single or low-copy genes across land plants, it is possible that paralogous or other non-targeted sequences were assembled from the enriched data. Thus, in step 4 (*orthology to reference*) we assessed orthology based on the best tblastx (Camacho et al. 2009) hit of the probe region of each assembled sequence to the coordinates of 10 plant genomes representing hornworts, liverworts, mosses, lycophytes, ferns, and gymnosperms (Supplementary Material). Since assemblies may extend into the flanking introns, we performed orthology assessment with the assembled probe regions. We called an assembled sequence an ortholog of the probe if it had no additional tblastx hits with >95% of the best bit score, outside of a 1000 base pair flanking window around the genomic coordinates of the probe locus in a reference genome. We only required that a sequence have evidence of orthology in one of the reference genomes. At this point in the pipeline, a taxon may retain more than one orthologous sequence for a single locus, potentially representing allelic variation or duplication.

In the fifth step (*contamination filter*), in order to filter out likely contaminants, for each assembled sequence we performed a tblastx search against the respective reference 1KP and genomic sequences for that locus. If a sequence’s best hit was not from the taxonomic group (i.e., hornwort, liverwort, moss, lycophyte, fern, or gymnosperm) from which the sequence came, that sequence was removed as a potential contaminant. Finally, in the sixth step (*alignment and merge isoforms*), we aligned the probe region-only sequences using MAFFT version 7.425 (Katoh and Standley 2013). Sequences from the same taxon with mismatches due to heterozygous sites were merged with a Perl script, using IUPAC codes to represent heterozygous sites.

In order to evaluate the usefulness of the probe set for phylogenetic inference, we ran a ML analysis on a supermatrix of the locus alignments. After completing the pipeline, it is possible that a sample would still have multiple sequences in an exon alignment where BRIDGER determined that reads represented more than simple allelic diversity, such as homeologs, paralogs, or alleles inherited though hybridization. In these cases, we retained the longest sequence and removed the other sequences from that sample. We also removed sequences from all samples from which we recovered fewer than 10% (i.e., 45) of the loci and then pruned the alignments so that they only included sites (i.e., columns) that had data from at least four samples. We concatenated all loci into a single supermatrix and ran a ML search and 100 nonparametric bootstrap (BS) replicates using RAxML 8.2.10 with the GTR CAT model (Stamatakis, 2014). The scripts used to process the data for phylogenetic analysis and the supermatrix alignment with locus boundaries are available on Dryad (https://doi.org/10.5061/dryad.7pvmcvdqg).

### Optimizing the GoFlag 451 Probe Set

Based on the results of this pilot study, we refined the original *GoFlag 451* probe set to optimize the performance of the target enrichment across flagellate plants. The resulting *GoFlag 408* probe set is a subset of the original *GoFlag 451* probe set, which contains 52,306 probes covering 408 of the original 451 loci. For the *GoFlag 408* probe set, we removed probes for all but two of the loci that produced sequences from fewer than 104 samples in this study, along with other probes that were either underperforming or exhibited strong taxonomic biases (Fig. 3; see Supplemental Table 1). The *GoFlag 408* probe set is also available on Dryad (https://doi.org/10.5061/dryad.7pvmcvdqg) and commercialized by RAPiD Genomics (http://rapid-genomics.com). Although we did not run a separate target enrichment experiment to assess the performance of the *GoFlag 408,* we examined the phylogenetic signal in the 408 selected loci based on data generated using the *GoFlag 451* probe set. Specifically, we made a concatenated matrix of just the probe regions corresponding to the *GoFlag 408* probe set, and we ran a ML phylogenetic analysis on that supermatrix as described above (data available on Dryad).

## RESULTS

The ML phylogenetic analysis of the supermatrix of the 451 exons used for the design of the probe set from the 1KP data provided a strongly supported land plant tree with relationships that are generally consistent with those from formal 1KP analyses (Supplemental Figure 1; One Thousand Plant Transcriptomes Initiative, 2019). Throughout the tree, 81.5% (767/941) of the internal branches had 100% BS support, 89.3% of the branches had at least 90% BS support, and 94.6% of the branches had at least 70% BS support (Supplemental Figure 1). This suggests that the 451 relatively conserved loci covered by the *GoFlag 451* probe set provide sufficient data to resolve many relationships throughout land plants, and in many cases appear to provide similar if not better resolution compared to the full 1KP single gene dataset (Supplemental Figure 1; One Thousand Plant Transcriptomes Initiative, 2019).

One measure of the performance of the probe set is the proportion of sequences from each library that mapped to the probe loci. The target enrichment ranged from 0.1% (*Aneura pinguis* (L.) Dumort, a liverwort) to 89.9% (*Rhynchostegium murale* (Hedw.) Schimp., a moss) of the reads, with an average across samples of 42.5% and a median of 40.9% (Supplemental Table 1). The number of loci recovered (out of a possible 451) ranged from 3 (*Mesoptychia badensis* (Gottsche ex Rabenh.) L.Söderstr. et Vána, a liverwort) to 436 (*Podocarpus smithii* de Laub, a gymnosperm), with an average of 332.4 and a median of 394.0 (Fig. 1; Supplemental Table 2). While we recovered fewer than 10% of the possible loci in 16 samples, in 82 of the 188 samples we recovered at least 90% of the possible loci (Fig. 1; Supplemental Table 2). Overall, the probes worked well across flagellate plant lineages, with the fewest average number of loci in the gymnosperm samples, and the most in the mosses (Table 1). There were 17 species in our target enrichment experiment that also had transcriptome data generated by 1KP (One Thousand Plant Transcriptomes Initiative, 2019). In 13 of the 17 common species, our target enrichment study generated data from more of the 451 loci than 1KP (Supplemental Table 3), suggesting either that these loci were missed in the transcriptome sequencing or our experiment amplified divergent copies that were excluded from the 1KP alignments.

**Figure 1.**
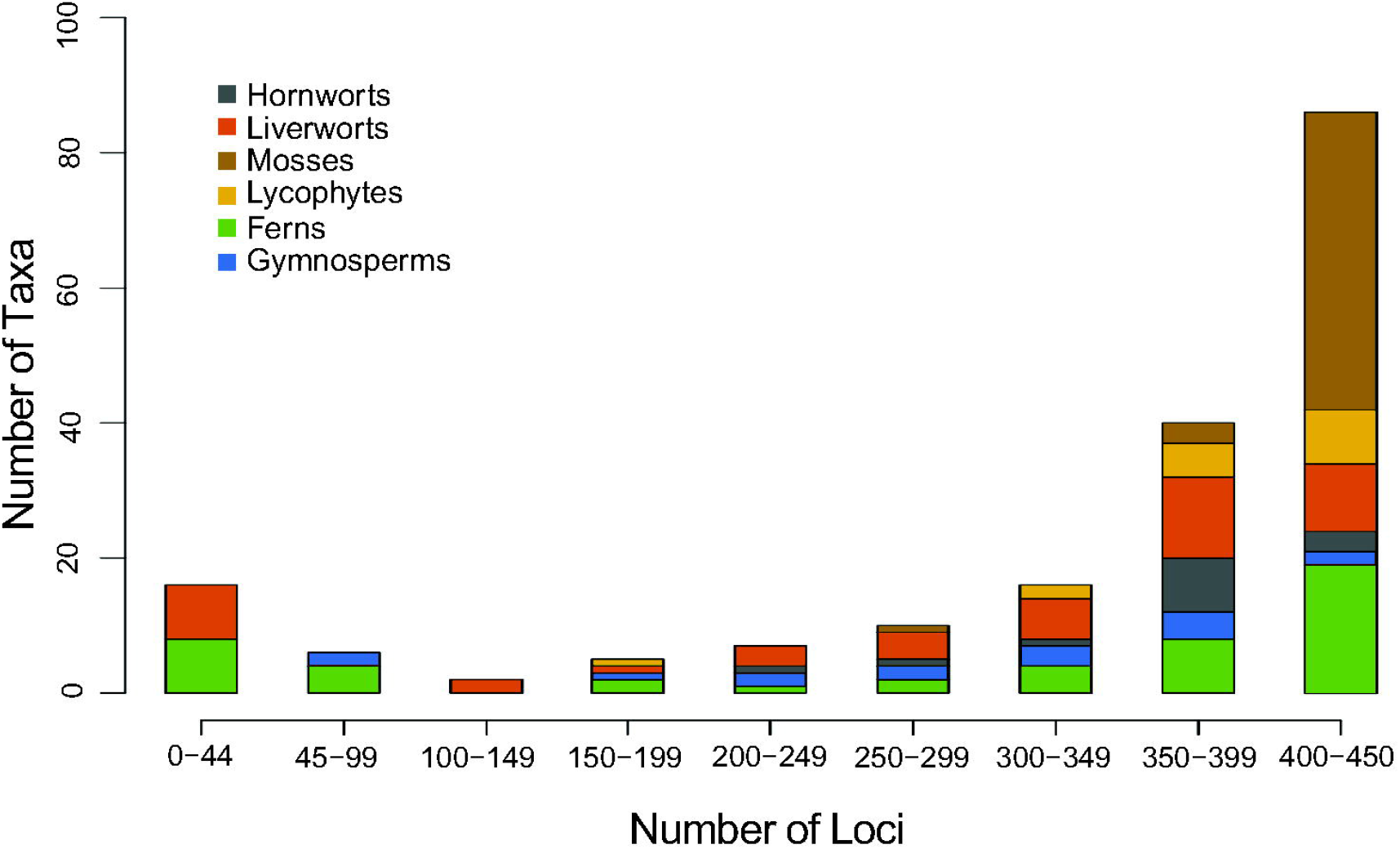
Distribution of number of loci successfully sequenced (out of a possible 451) per taxon sample. Colors represent the lineages of the samples. Each locus is a relatively conserved exon from a single or low-copy nuclear gene.

**Table 1.**
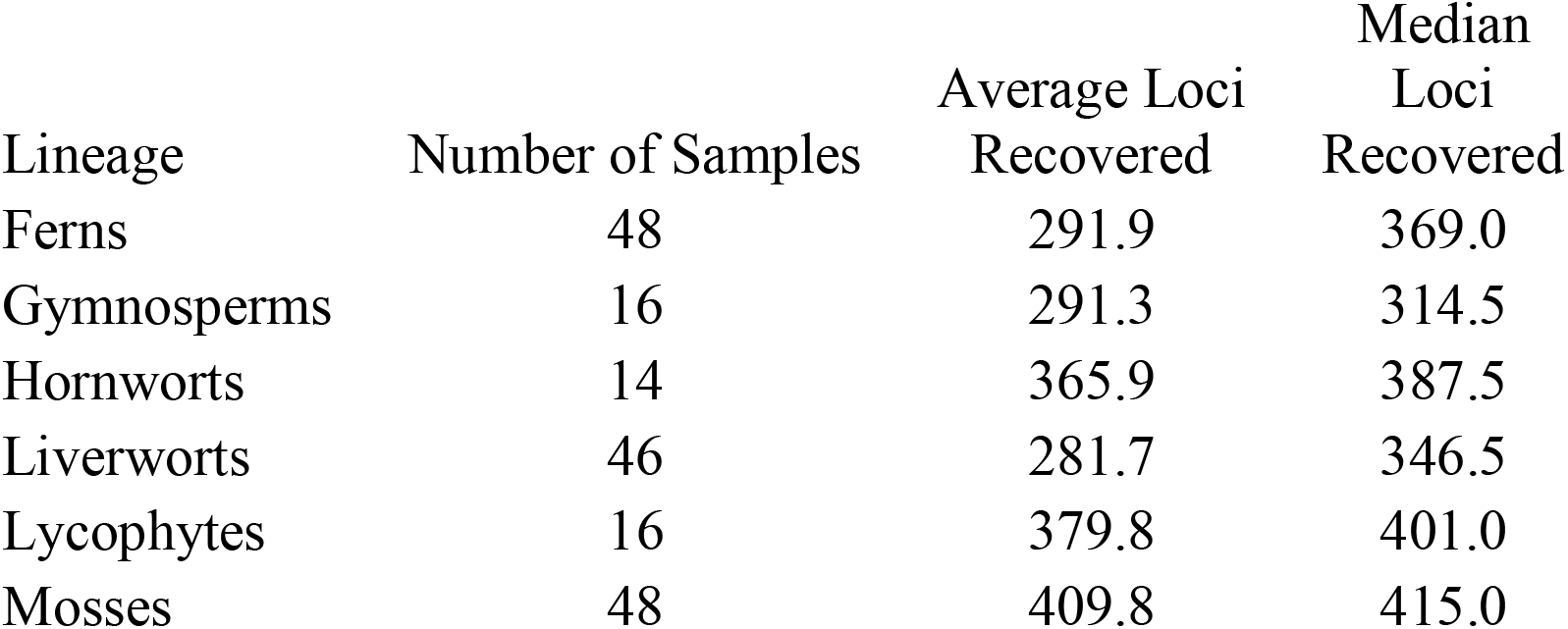
Distribution of loci with sequence data (out of a possible 451) across major flagellate plant lineages.

| Lineage | Number of Samples | Average Loci<br>Recovered | Median<br>Loci<br>Recovered |
| --- | --- | --- | --- |
| Ferns | 48 | 291.9 | 369.0 |
| Gymnosperms | 16 | 291.3 | 314.5 |
| Hornworts | 14 | 365.9 | 387.5 |
| Liverworts | 46 | 281.7 | 346.5 |
| Lycophytes | 16 | 379.8 | 401.0 |
| Mosses | 48 | 409.8 | 415.0 |

The samples for which we recovered few loci could have had highly diverged sequences from the probe sites or had poor quality DNA. However, in a few cases species from which we recovered few loci are closely related to species from which we recovered many loci (e.g., *Dryopteris pentheri* (Krasser) C. Chr., 21 loci, vs. *Dryopteris patula* (Sw.) Underw, 431 loci, or *Elaphoglossum yatesii* (Sodiro) Christ, 27 loci, vs. *Elaphoglossum bellermannianum* (Klotzsch) T. Moore, 418 loci; Supplemental Table 2), suggesting that probe site evolution is unlikely to explain at least some of the failed captures. Similarly, we found no relationship between the amount of DNA from a given specimen and the number of recovered loci (Fig. 2A; although note that the input DNA into the library was normalized to 250 ng, meaning that we did not use more than 250 ng of DNA for any samples, even if they had more than 250 ng of DNA). Some samples with very little DNA were successful, and some samples with abundant DNA were not (Fig. 2A), suggesting that DNA quality rather than quantity may be affecting these libraries. However, the samples from which we recovered few loci all had relatively few reads (Fig. 2B). We obtained sequence data from an average of 138.6 (median = 147.0) out of 188 total samples across the 451 loci (Supplemental Table 1), but there also was variation in the number of samples that recovered each locus, and some loci had a taxonomic bias (Fig. 3; Supplemental Table 1).

**Figure 2.**
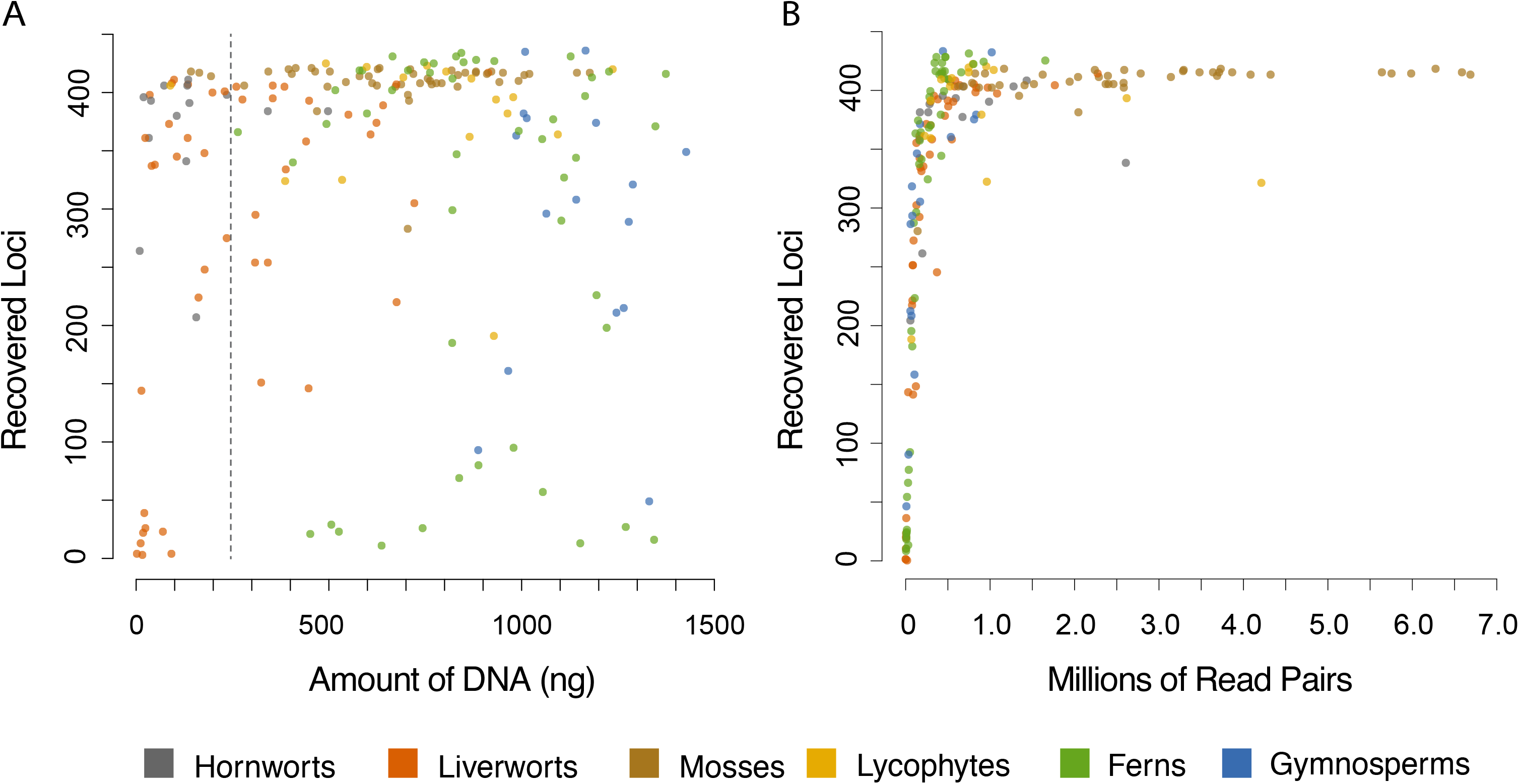
A) Amount of DNA in each sample vs. the number of resulting loci obtained in the targeted enrichment analysis. The dashed line at 250 ng represents the amount of DNA at which samples were normalized for the library preparation. B) Number of reads obtained from each sample vs. the number of loci obtained from the targeted enrichment analysis. Colors represent the major lineage of the sample.

**Figure 3.**
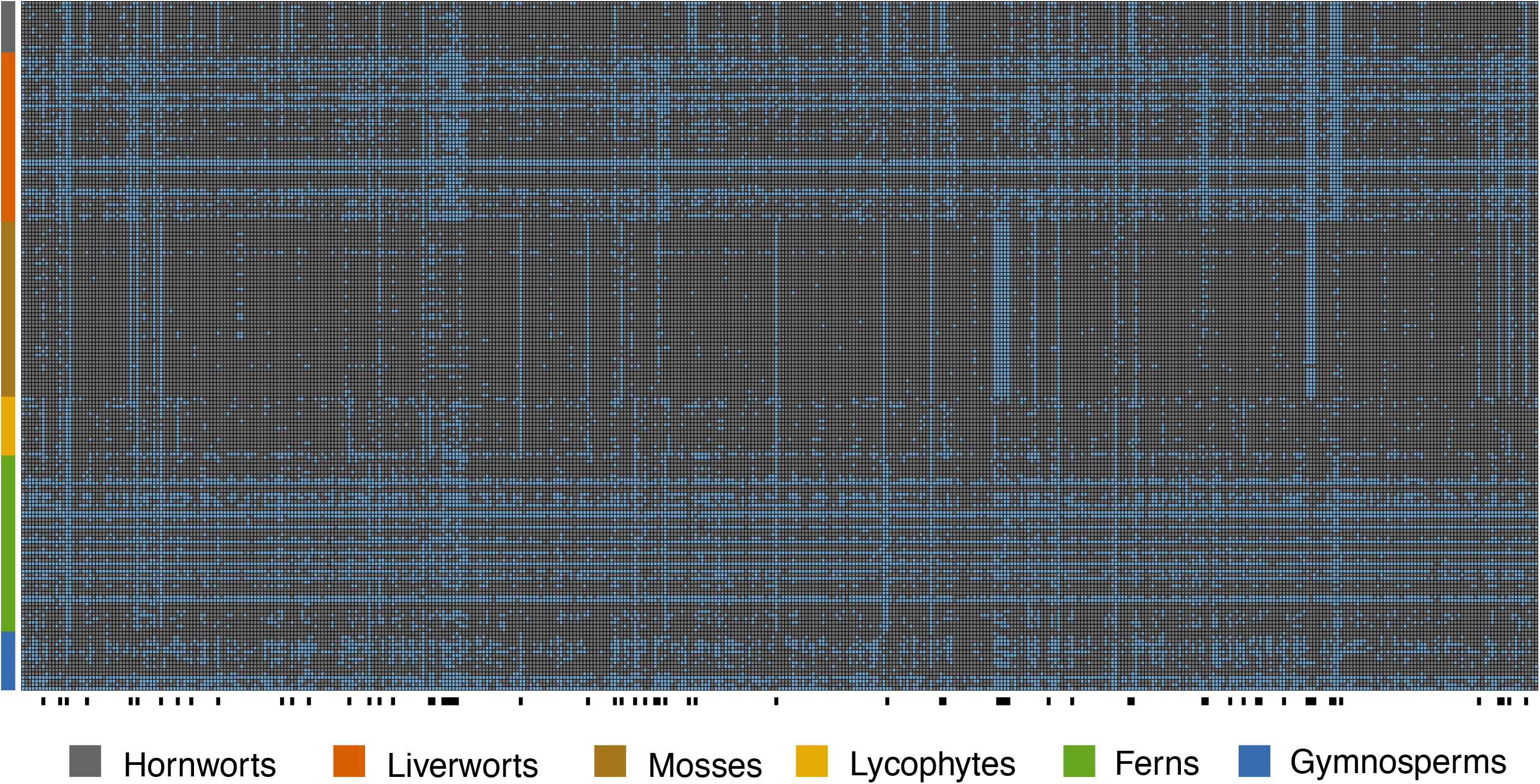
Heat map showing the distribution of data in the flagellate plant samples across the 451 probe regions (i.e., exons). Loci that were missing for an individual are colored blue in the heatmap while sampled loci are grey. Black bars along the bottom of the heatmap indicate loci with biases among major plant groups, where less than 25% of one group had the locus and over 75% of another group had the locus.

To evaluate the phylogenetic signal in the data, we constructed a 172-taxon phylogenetic supermatrix of the 451 loci (i.e., exonic probe regions) that was 90,153 nucleotides in length and 75.5% full (i.e., 24.5% missing data). Of the 170 clades in the ML tree, 139 (82%) had 100% BS support; 90% of the clades had at least 90% BS support, and only 5 clades had less than 70% BS support (Fig. 4). The resulting phylogenetic tree is generally consistent with the consensus land plant phylogeny (e.g., One Thousand Plant Transcriptomes Initiative, 2019). One unexpected result is the non-monophyly of the two *Targionia hypophylla* L. (liverwort) samples (Fig. 4). This could be the result of misidentification of the specimes; however, many of the bryophytes were sampled from mixed herbarium samples that contained tissue from multiple taxa. This may also explain the relatively large number of contaminant sequences identified in many of the bryophytes (Supplemental Table 1).

**Figure 4.**
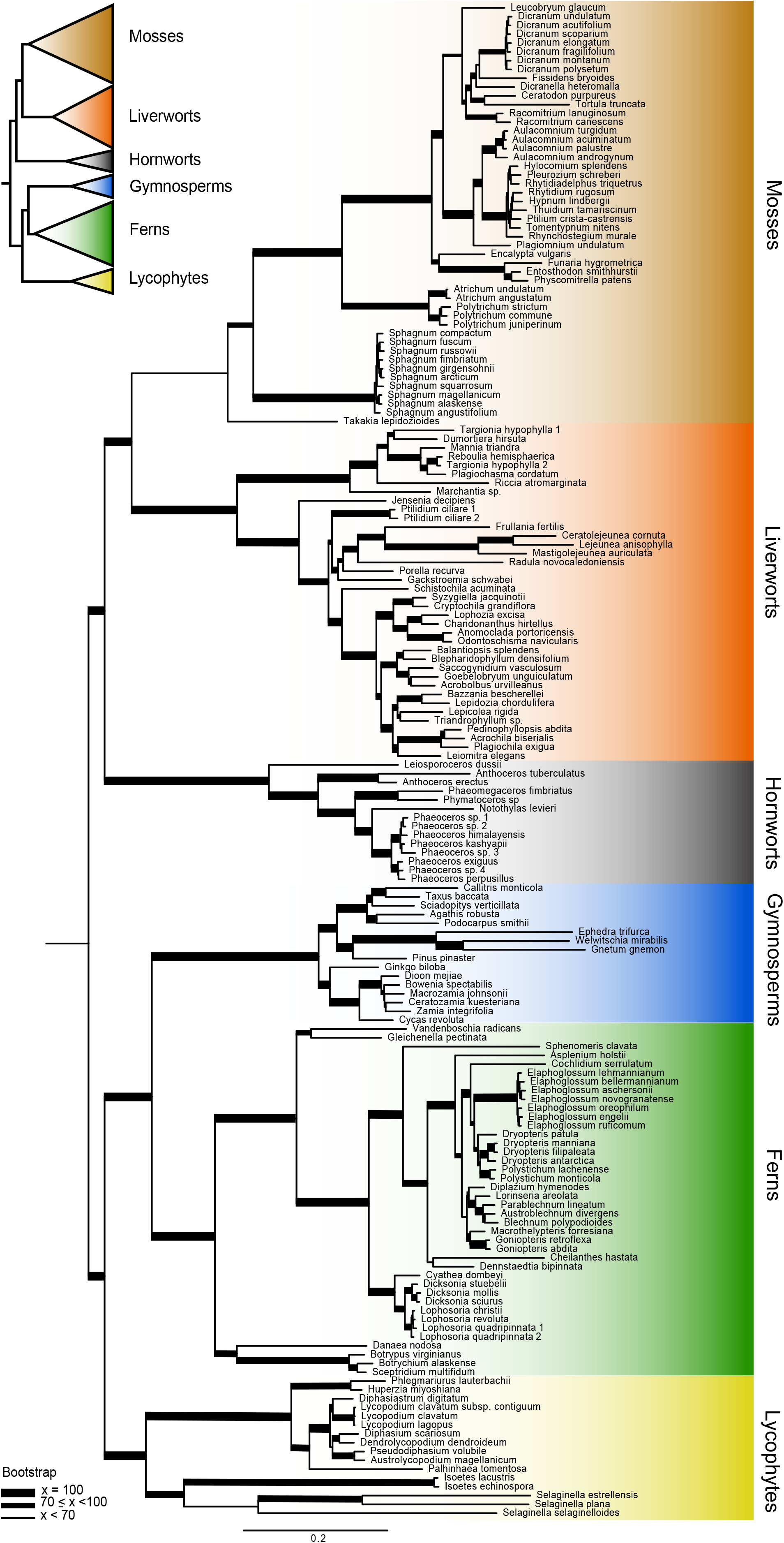
Phylogram from a ML analysis of the supermatrix made by concatenating the alignments from the *GoFlag 451* probe regions (i.e., exons) for the samples with at least 45 loci. The tree was arbitrarily rooted between the bryophytes and vascular plants.

To explore the potential for the probe set to resolve relationships among closely related taxa, we assembled supermatrices including both the probe regions (i.e., conserved exons) and the more variable flanking regions for samples from the seven genera from which we had at least four samples. In the supermatrices we only included loci with data from at least four taxa, and within each locus alignment, we only included columns with at least four nucleotides. By including the flanking regions, the length of the alignments was between 1.8 and 6.0 times longer than the probe region-only alignments, with between 2.5 and 10.7 times more variable sites (i.e., columns in the alignment that have at least two different nucleotides; Table 2). In contrast to the exonic probe regions, which can be easily aligned across land plants, it can be difficult to align the variable flanking regions across distantly related taxa. Nevertheless, the flanking regions potentially can provide a tremendous amount of additional data to infer phylogenies among more closely related taxa.

**Table 2.** Comparison of phylogenetic data from probe regions (i.e., exons) and the probe + flanking regions for the seven genera with at least four samples.

| Genus | Lineage | Samples | Loci | Probe Region Only |  | Probe & Flanking Regions |  |
| --- | --- | --- | --- | --- | --- | --- | --- |
|  |  |  |  | Alignment<br>(bp) | Variable<br>Characters | Alignment<br>(bp) | Variable<br>Characters |
| *Aulacomnium* | Moss | 4 | 414 | 75492 | 2846 | 422123 | 30542 |
| *Dicranum* | Moss | 7 | 415 | 76177 | 2341 | 441413 | 19510 |
| *Dryopteris* | Fern | 5 | 223 | 48437 | 3158 | 86463 | 7955 |
| *Elaphoglossum* | Fern | 8 | 386 | 74673 | 2165 | 185268 | 8118 |
| *Lophosoria* | Fern | 4 | 417 | 78941 | 1014 | 223274 | 4250 |
| *Phaeoceros* | Hornwort | 8 | 397 | 75394 | 5803 | 252294 | 22148 |
| *Sphagnum* | Moss | 10 | 409 | 74202 | 3951 | 444271 | 39242 |

Finally, the supermatrix for the loci in the optimized *GoFlag 408* probe set from the samples with data from at least 45 loci was 80,748 bp long and 79.9% full. Although this supermatrix alignment was 9,405 bp shorter than the supermatrix made from the *GoFlag 451* loci, the topology and levels of support from the resulting trees were virtually identical (Supplemental Figure 2).

## CONCLUSIONS

Here we have described a probe set targeting nuclear loci across flagellate plants that diverged as much as ~450 million years ago. The probe region (i.e., exon) sequences are easily aligned across land plants and can help resolve backbone phylogenetic relationships among flagellate plant lineages (Fig. 4; Supplemental Figures 1,2). Furthermore, the more variable flanking regions provide abundant data for resolving relationships among closely related species, or potentially even populations within a species (Table 2). Although the *GoFlag 451* probe set worked well in all major extant flagellate plant lineages (Table 1; Figs. 1, 3), the sampling from this study is not sufficient to determine if the probe set will work well in all flagellate plant taxa. Our strategy was to develop a “universal” probe set that covers the majority of these groups, and the *GoFlag 451* probe set and the analysis pipeline provide a core set of validated tools accessible to all scientists. However, some evolutionary questions in the flagellate plants may require a more specific probe set for more closely related taxa (e.g., Larridon et al., 2020). While the *GoFlag 451* probe set facilitates target enrichment projects in any flagellate plant group, a probe set designed for a particular lineage could easily have more specific probes that cover either more loci, loci of special interest (e.g., Medina et al., 2019; Montes et al., 2019), or loci with higher substitution rates (de La Harpe et al. 2019). Resolving some of the more contentious flagellate plant relationships may likewise require a larger, more specific probe set. In those cases, the *GoFlag 451* probes define a core set of loci that can be built upon. Nuclear gene evolution within land plants is often extremely complex, with, for example, frequent gene and whole genome duplications. Although nuclear loci have the potential to resolve complex evolutionary relationships, their own complex histories can easily mislead and complicate phylogenetic inference. Our test for orthology in the analytical pipeline is simplistic, and in this study, we did not carefully examined potential issues of paralogy or homoeology in the 451 loci within flagellate plants. However, the resulting sequence data can be used to examine gene or genome duplication, or even allelic variation and heterozygosity.

In subsequent sequencing runs, the GoFlag project has used the *GoFlag 408* probe set, and results indicate similar, if not better, overall performance compared to the *GoFlag 451* probe set (JGB, unpublished observation). Due to the large number of probes needed to cover the diversity of flagellate plants, we did not include the angiosperms when designing the GoFlag probe sets. However, the same exons appear to be conserved across angiosperms and provide sufficient data to resolve many angiosperm relationships (Supplemental Figure 1). Thus, the loci in this probe set may provide a foundation for constructing large-scale nuclear phylogenies across land plants.

## Acknowledgements

This work was funded by the U.S. National Science Foundation (DEB-1541506). We thank Jim Leebens-Mack and Gane Wong for early access to 1KP transcriptome data, Matt Johnson for discussions and advice about probe design, and Adam Payton for lab help, especially with scaling up the DNA extraction capacity.

## Data Availability

Sequence reads have been deposited to the National Center for Biotechnology (NCBI) Sequence Read Archive (PRJNA630729). The *GoFlag 451* and *GoFlag 408* probe sets are available on Dryad (https://doi.org/10.5061/dryad.7pvmcvdqg) with the pipeline scripts and refence sequences, the post-processing scripts, and all phylogenetic matrices and trees from this study. Accessions and voucher information are in Supplemental Table 2.

---

Supplemental Figure 1. Phylogram from an ML analysis of a supermatrix made by concatenating 1KP transcriptome sequences from the gene regions covered by the *GoFlag 451* probe set. The tree was arbitrarily rooted between the bryophytes and vascular plants.

Supplemental Figure 2. Phylogram from a ML analysis of the supermatrix made by concatenating the alignments from the *GoFlag 408* probe regions (i.e., exons) for the samples with at least 45 loci. The tree was arbitrarily rooted between the bryophytes and vascular plants.

