## Supplemental Table 1 for "A target enrichment probe set for resolving the flagellate plant tree of life"

**Supplemental Table 1**. Description of the loci from the GoFlag 451 probe set. "Total Reads" is the number of reads from all samples that map to the locus. "Samples" is the number of samples that recovered the locus in the targeted enrichment experiment. The "Cut in GoFlag 408 Probe Set" denotes the loci that were not included in the Go Flag 408 probe set, and a '*' identifies the loci with taxonomic biases in the samples that recovered the locus. "1KP Locus" identifies the 1KP locus and exon region in which the probe region is found.

| **Locus** | **Total Reads** | **Samples** | **Ave. Probe Region Lenth (bp)** | **Cut in GoFlag 408 Probe Set** | **1KP Locus** |
| --- | --- | --- | --- | --- | --- |
| 1 | 345242 | 131 | 178.0 | No | L4471_0 |
| 2 | 232182 | 129 | 138.8 | No | L4519_1 |
| 3 | 333396 | 143 | 119.1 | No | L4519_3 |
| 4 | 320547 | 143 | 131.8 | No | L4519_6 |
| 5 | 347704 | 130 | 131.9 | No | L4519_7 |
| 6 | 315847 | 131 | 133.5 | No | L4519_8 |
| 7 | 803488 | 154 | 347.2 | No | L4527_0 |
| 8 | 499663 | 157 | 166.9 | No | L4603_1 |
| 9 | 817557 | 157 | 328.3 | No | L4603_2 |
| 10 | 667084 | 165 | 238.7 | No | L4603_4 |
| 11 | 752448 | 163 | 242.3 | No | L4691_0 |
| 12 | 516353 | 169 | 241.6 | No | L4724_0 |
| 13 | 29575 | 13 | 423.5 | No* | L4744_0 |
| 14 | 445531 | 145 | 177.1 | No | L4744_3 |
| 15 | 506243 | 157 | 148.9 | No | L4757_1 |
| 16 | 476627 | 149 | 171.3 | No | L4757_2 |
| 17 | 609538 | 163 | 156.0 | No | L4793_3 |
| 18 | 769580 | 156 | 245.6 | No | L4793_7 |
| 19 | 428698 | 149 | 177.6 | No | L4796_1 |
| 20 | 646566 | 163 | 223.2 | No | L4802_0 |
| 21 | 298313 | 151 | 223.8 | No | L4806_0 |
| 23 | 651466 | 164 | 259.5 | No | L4848_1 |
| 24 | 1069064 | 169 | 282.8 | No | L4848_5 |
| 25 | 321200 | 148 | 145.6 | No | L4889_1 |
| 26 | 336581 | 150 | 118.7 | No | L4889_2 |
| 27 | 292417 | 138 | 151.3 | No* | L4889_3 |
| 28 | 247938 | 140 | 137.0 | No | L4890_0 |
| 29 | 301034 | 146 | 143.8 | No | L4893_0 |
| 30 | 302772 | 150 | 140.2 | No | L4893_1 |
| 31 | 1055685 | 160 | 449.4 | No | L4932_0 |
| 32 | 932487 | 149 | 167.0 | No | L4932_1 |
| 33 | 791928 | 164 | 221.4 | No | L4942_0 |
| 34 | 355864 | 156 | 161.8 | No | L4942_1 |
| 35 | 856084 | 150 | 254.8 | No | L4951_0 |
| 36 | 975191 | 165 | 203.3 | No | L4951_1 |
| 37 | 129900 | 101 | 139.9 | No | L4976_2 |
| 38 | 377808 | 152 | 119.5 | No | L4989_1 |
| 39 | 413538 | 160 | 147.1 | No | L4989_3 |
| 40 | 369787 | 138 | 157.1 | No | L4992_1 |
| 41 | 223936 | 123 | 143.1 | No | L4992_12 |
| 42 | 217629 | 127 | 140.6 | No | L4992_13 |
| 43 | 332156 | 141 | 143.8 | No | L4992_3 |
| 44 | 356221 | 157 | 175.6 | No | L4992_4 |
| 45 | 413911 | 149 | 129.8 | No | L4992_5 |
| 46 | 402686 | 152 | 169.6 | No | L4992_7 |
| 47 | 354780 | 152 | 197.3 | No | L4992_9 |
| 48 | 419808 | 151 | 129.7 | No | L5018_0 |
| 49 | 567093 | 153 | 184.8 | No | L5018_1 |
| 50 | 515388 | 154 | 143.3 | No | L5018_3 |
| 51 | 441549 | 138 | 159.0 | No | L5018_4 |
| 52 | 493807 | 162 | 164.4 | No | L5018_5 |
| 53 | 296309 | 149 | 138.0 | No | L5032_1 |
| 54 | 850964 | 153 | 166.5 | No | L5034_1 |
| 55 | 452246 | 149 | 127.3 | No | L5090_1 |
| 56 | 304421 | 147 | 154.1 | No | L5111_0 |
| 57 | 560674 | 145 | 221.2 | No | L5116_1 |
| 58 | 465841 | 145 | 171.8 | No | L5116_3 |
| 59 | 617108 | 166 | 191.5 | No | L5138_3 |
| 60 | 690477 | 156 | 193.5 | No | L5163_0 |
| 61 | 398225 | 167 | 177.1 | No | L5163_2 |
| 62 | 309352 | 131 | 137.6 | No | L5163_3 |
| 63 | 443091 | 145 | 149.2 | No | L5163_5 |
| 64 | 366729 | 156 | 119.8 | No | L5163_6 |
| 65 | 305326 | 150 | 128.6 | No | L5163_7 |
| 66 | 452608 | 143 | 122.5 | No | L5168_1 |
| 67 | 416720 | 127 | 146.6 | No | L5168_2 |
| 68 | 405161 | 151 | 214.1 | No | L5168_3 |
| 69 | 568946 | 153 | 192.7 | No | L5177_1 |
| 70 | 730924 | 164 | 170.5 | No | L5188_0 |
| 71 | 391939 | 152 | 184.7 | No | L5188_2 |
| 72 | 847671 | 163 | 209.8 | No | L5200_1 |
| 73 | 368782 | 145 | 152.7 | No | L5206_1 |
| 74 | 356770 | 151 | 131.6 | No | L5257_0 |
| 75 | 535763 | 156 | 197.5 | No | L5257_1 |
| 76 | 549387 | 150 | 167.3 | No | L5257_3 |
| 77 | 483414 | 159 | 211.1 | No | L5257_4 |
| 78 | 395301 | 154 | 125.3 | No | L5264_0 |
| 79 | 724927 | 161 | 252.6 | No | L5264_1 |
| 80 | 359151 | 141 | 133.0 | No | L5271_0 |
| 81 | 415344 | 157 | 199.8 | No | L5271_2 |
| 82 | 553144 | 92 | 611.4 | CUT* | L5273_3 |
| 83 | 1988309 | 172 | 484.8 | No | L5280_1 |
| 84 | 325010 | 149 | 125.7 | No | L5296_1 |
| 85 | 284329 | 145 | 152.5 | No | L5296_2 |
| 86 | 294029 | 119 | 163.7 | No | L5299_1 |
| 87 | 439985 | 144 | 123.8 | No | L5318_0 |
| 88 | 223114 | 85 | 384.9 | CUT* | L5328_0 |
| 89 | 999958 | 125 | 420.9 | No | L5333_1a |
| 90 | 189313 | 102 | 346.8 | No* | L5333_1c |
| 91 | 371226 | 147 | 137.1 | No | L5333_1d |
| 93 | 237819 | 135 | 158.1 | No | L5335_2 |
| 94 | 199257 | 136 | 122.2 | No | L5339_0 |
| 95 | 327303 | 137 | 151.8 | No | L5339_1 |
| 96 | 443527 | 74 | 493.4 | CUT* | L5343_0 |
| 97 | 493355 | 158 | 154.8 | No | L5355_0 |
| 98 | 739843 | 164 | 264.3 | No | L5355_1 |
| 99 | 838012 | 165 | 167.8 | No | L5357_0 |
| 100 | 250771 | 136 | 122.6 | No | L5366_0 |
| 101 | 333715 | 139 | 122.8 | No | L5366_1 |
| 102 | 134351 | 113 | 135.0 | No | L5398_0 |
| 103 | 474414 | 154 | 195.5 | No | L5398_1 |
| 104 | 82270 | 97 | 131.0 | CUT* | L5404_2 |
| 105 | 190521 | 138 | 127.7 | No | L5404_4 |
| 106 | 730155 | 160 | 190.4 | No | L5406_1 |
| 107 | 798910 | 165 | 350.6 | No | L5406_3 |
| 108 | 670872 | 114 | 178.2 | No | L5406_5 |
| 109 | 509666 | 81 | 122.3 | CUT* | L5406_6 |
| 110 | 231200 | 50 | 125.9 | CUT* | L5421_1 |
| 111 | 4993 | 10 | 134.5 | CUT | L5426_0 |
| 112 | 1258439 | 163 | 319.4 | No | L5426_1 |
| 113 | 1447725 | 168 | 600.8 | No | L5426_3 |
| 114 | 852590 | 159 | 212.2 | No | L5426_5 |
| 115 | 557751 | 150 | 174.6 | No | L5426_6 |
| 116 | 309624 | 130 | 181.9 | No* | L5427_1 |
| 117 | 293260 | 126 | 132.0 | No | L5430_1 |
| 118 | 848920 | 160 | 391.2 | No | L5449_1 |
| 119 | 656039 | 157 | 228.9 | No | L5449_3 |
| 120 | 498207 | 159 | 204.6 | No | L5449_4 |
| 121 | 214883 | 122 | 148.7 | No | L5463_0 |
| 122 | 297498 | 145 | 152.6 | No | L5463_2 |
| 123 | 507937 | 155 | 185.6 | No | L5464_0 |
| 124 | 525129 | 146 | 125.5 | No | L5464_1 |
| 125 | 693921 | 158 | 255.1 | No | L5469_0 |
| 126 | 475412 | 155 | 148.8 | No | L5469_1 |
| 127 | 199305 | 126 | 128.2 | No | L5477_1 |
| 128 | 103462 | 79 | 177.9 | CUT* | L5502_0 |
| 129 | 222923 | 116 | 138.9 | No | L5513_1 |
| 130 | 1417498 | 153 | 140.1 | No | L5528_0 |
| 131 | 424689 | 127 | 120.4 | No | L5528_1 |
| 132 | 244550 | 142 | 140.6 | No | L5528_3 |
| 133 | 538101 | 150 | 152.0 | No | L5528_4 |
| 134 | 59874 | 28 | 222.3 | CUT | L5536_0 |
| 135 | 475547 | 148 | 128.8 | No | L5551_2 |
| 136 | 40529 | 39 | 211.0 | CUT* | L5554_0 |
| 137 | 242586 | 148 | 139.9 | No | L5554_1 |
| 138 | 470523 | 157 | 187.7 | No | L5554_4 |
| 139 | 247035 | 145 | 136.7 | No | L5554_6 |
| 140 | 247510 | 119 | 193.1 | No* | L5596_0 |
| 141 | 672931 | 147 | 247.7 | No | L5596_6 |
| 142 | 435116 | 149 | 156.9 | No | L5599_0 |
| 143 | 404406 | 146 | 200.6 | No | L5599_1 |
| 144 | 435143 | 143 | 124.9 | No* | L5599_4 |
| 145 | 600966 | 157 | 213.3 | No | L5614_0 |
| 146 | 588573 | 170 | 282.1 | No | L5614_1 |
| 147 | 454786 | 146 | 137.5 | No | L5614_3 |
| 148 | 217871 | 134 | 129.9 | No | L5634_0 |
| 149 | 508528 | 152 | 128.9 | No | L5639_0 |
| 150 | 532447 | 156 | 165.2 | No | L5639_1 |
| 151 | 162724 | 119 | 125.3 | No* | L5644_0 |
| 152 | 765843 | 164 | 178.1 | No | L5644_1 |
| 153 | 768914 | 165 | 246.6 | No | L5644_2 |
| 154 | 238392 | 142 | 143.8 | No | L5644_3 |
| 155 | 467644 | 146 | 211.6 | No | L5664_0 |
| 156 | 523531 | 146 | 134.6 | No | L5664_1 |
| 157 | 517539 | 149 | 140.5 | No | L5670_1 |
| 158 | 519063 | 153 | 221.0 | No | L5670_3 |
| 159 | 419481 | 158 | 147.5 | No | L5702_1 |
| 160 | 434490 | 160 | 173.7 | No | L5702_3 |
| 161 | 493961 | 147 | 172.7 | No | L5703_1 |
| 162 | 558709 | 166 | 173.1 | No | L5703_4 |
| 163 | 691248 | 163 | 202.4 | No | L5721_1 |
| 164 | 289295 | 128 | 182.6 | No | L5753_6 |
| 165 | 439592 | 151 | 176.4 | No | L5761_0 |
| 166 | 236394 | 140 | 128.5 | No | L5770_0 |
| 167 | 383521 | 157 | 184.1 | No | L5770_1 |
| 168 | 165263 | 127 | 149.6 | No | L5770_3 |
| 169 | 387180 | 116 | 128.7 | No* | L5802_0 |
| 170 | 553047 | 152 | 190.9 | No | L5812_2 |
| 171 | 416075 | 141 | 125.7 | No* | L5815_0 |
| 172 | 552743 | 154 | 170.4 | No | L5815_1 |
| 173 | 290969 | 135 | 128.9 | No | L5816_0 |
| 174 | 250593 | 141 | 141.2 | No | L5816_1 |
| 175 | 369263 | 147 | 167.2 | No | L5818_1 |
| 176 | 289289 | 119 | 129.1 | No* | L5818_3 |
| 177 | 563354 | 153 | 149.4 | No | L5821_0 |
| 178 | 579812 | 146 | 119.7 | No | L5821_1 |
| 179 | 781275 | 167 | 226.8 | No | L5821_2 |
| 180 | 577130 | 158 | 173.5 | No | L5821_3 |
| 181 | 582922 | 156 | 167.0 | No | L5822_0 |
| 182 | 340312 | 131 | 147.5 | No | L5822_1 |
| 183 | 733764 | 169 | 252.0 | No | L5840_0 |
| 184 | 658157 | 155 | 194.6 | No | L5840_1 |
| 185 | 636343 | 142 | 173.0 | No | L5840_3 |
| 186 | 255739 | 133 | 130.5 | No | L5841_0 |
| 187 | 139890 | 107 | 137.7 | No* | L5842_0 |
| 188 | 249397 | 134 | 161.0 | No | L5842_1 |
| 189 | 285080 | 125 | 126.2 | No | L5842_4 |
| 190 | 280641 | 140 | 128.2 | No | L5842_5 |
| 191 | 569173 | 149 | 145.6 | No | L5849_0 |
| 192 | 310579 | 76 | 126.0 | CUT* | L5849_2 |
| 193 | 685016 | 138 | 260.4 | No | L5853_0 |
| 194 | 665049 | 163 | 282.2 | No | L5857_1b |
| 195 | 1012521 | 54 | 206.3 | CUT* | L5858_0 |
| 196 | 323135 | 143 | 157.5 | No | L5863_4 |
| 197 | 823856 | 157 | 297.3 | No | L5865_1 |
| 198 | 445904 | 147 | 212.0 | No | L5866_0 |
| 199 | 396707 | 119 | 121.1 | No* | L5870_0 |
| 200 | 329457 | 139 | 122.6 | No | L5892_1 |
| 201 | 393906 | 149 | 131.6 | No | L5894_1 |
| 202 | 1057483 | 160 | 375.2 | No | L5899_0 |
| 203 | 324664 | 124 | 158.5 | No | L5899_1 |
| 204 | 286001 | 153 | 135.9 | No | L5899_6 |
| 205 | 764096 | 168 | 417.4 | CUT | L5910_0 |
| 206 | 272527 | 43 | 312.3 | CUT | L5910_0b |
| 207 | 1436619 | 172 | 449.6 | No | L5913_1 |
| 208 | 130467 | 77 | 174.8 | CUT* | L5921_0 |
| 209 | 190913 | 94 | 158.0 | CUT* | L5921_1 |
| 210 | 257119 | 139 | 296.4 | No | L5921_10 |
| 211 | 188575 | 88 | 175.7 | CUT* | L5921_11 |
| 212 | 149385 | 102 | 208.6 | No* | L5921_12 |
| 213 | 129024 | 95 | 119.7 | CUT* | L5921_13 |
| 214 | 119996 | 89 | 148.9 | No* | L5921_14 |
| 215 | 313432 | 103 | 134.9 | No* | L5921_15 |
| 216 | 37245 | 59 | 126.9 | CUT | L5921_3 |
| 217 | 146466 | 118 | 185.2 | No | L5921_7 |
| 218 | 187687 | 118 | 131.6 | No | L5921_8 |
| 219 | 333845 | 157 | 154.8 | No | L5922_1 |
| 220 | 364627 | 161 | 191.0 | No | L5922_4 |
| 221 | 323401 | 147 | 149.5 | No | L5922_5 |
| 222 | 307368 | 145 | 152.6 | No | L5922_6 |
| 223 | 267194 | 147 | 135.0 | No | L5926_0 |
| 224 | 273488 | 130 | 125.7 | No | L5926_2 |
| 225 | 414412 | 152 | 140.1 | No | L5933_1 |
| 226 | 566785 | 171 | 273.5 | No | L5933_3 |
| 227 | 346220 | 149 | 171.8 | No | L5933_4 |
| 228 | 573181 | 148 | 146.0 | No | L5941_0 |
| 229 | 267324 | 148 | 137.9 | No | L5943_1 |
| 230 | 381276 | 154 | 184.0 | No | L5943_4 |
| 231 | 378632 | 145 | 129.0 | No | L5943_5 |
| 232 | 495760 | 159 | 146.4 | No | L5944_0 |
| 233 | 73700 | 67 | 134.0 | CUT* | L5944_1 |
| 234 | 680720 | 157 | 222.0 | No | L5944_3 |
| 235 | 1247950 | 164 | 232.7 | No | L5945_1 |
| 236 | 595628 | 156 | 175.4 | No | L5949_0 |
| 237 | 444072 | 159 | 119.5 | No | L5949_1 |
| 238 | 550302 | 145 | 132.1 | No | L5950_0 |
| 239 | 632767 | 160 | 173.6 | No | L5950_1 |
| 240 | 1247516 | 169 | 361.2 | No | L5958_0 |
| 241 | 399498 | 139 | 192.3 | No | L5960_0 |
| 242 | 426039 | 158 | 197.0 | No | L5960_1 |
| 243 | 592664 | 144 | 190.8 | No | L5974_2 |
| 244 | 435308 | 134 | 125.6 | No | L5980_1 |
| 245 | 601341 | 158 | 207.6 | No | L5981_1 |
| 246 | 467058 | 156 | 155.7 | No | L6000_0 |
| 247 | 513225 | 139 | 128.6 | No | L6000_2 |
| 248 | 929678 | 176 | 278.7 | No | L6016_1 |
| 249 | 324180 | 153 | 122.6 | No | L6026_1 |
| 250 | 515698 | 145 | 173.6 | No | L6034_2 |
| 251 | 213238 | 92 | 214.5 | CUT* | L6035_5 |
| 252 | 1860912 | 148 | 478.7 | No | L6035_6 |
| 253 | 917692 | 164 | 262.1 | No | L6036_1 |
| 254 | 871220 | 154 | 176.0 | No | L6036_5 |
| 255 | 742510 | 166 | 166.8 | No | L6038_1 |
| 256 | 572187 | 151 | 261.1 | No | L6041_1 |
| 257 | 518672 | 166 | 215.9 | No | L6041_6 |
| 258 | 401185 | 133 | 152.5 | No | L6041_7 |
| 259 | 274103 | 82 | 121.3 | CUT* | L6041_8 |
| 260 | 183652 | 110 | 265.6 | No* | L6051_0b |
| 261 | 286966 | 143 | 151.7 | No | L6056_2 |
| 262 | 305732 | 147 | 159.2 | No | L6056_4 |
| 263 | 618053 | 129 | 213.6 | No | L6064_0 |
| 264 | 184901 | 87 | 153.1 | CUT* | L6065_0 |
| 265 | 232837 | 129 | 155.5 | No | L6065_1 |
| 266 | 277022 | 157 | 205.4 | No | L6098_0 |
| 267 | 696877 | 113 | 140.7 | No* | L6098_1 |
| 268 | 743234 | 168 | 280.2 | No | L6098_4 |
| 269 | 719799 | 160 | 281.1 | No | L6098_5 |
| 270 | 593222 | 99 | 748.0 | No* | L6119_0 |
| 271 | 335328 | 137 | 168.7 | No | L6119_1 |
| 272 | 112056 | 81 | 123.8 | CUT* | L6119_3 |
| 273 | 123414 | 114 | 126.7 | No | L6119_4 |
| 274 | 311586 | 156 | 146.7 | No | L6130_1 |
| 275 | 952470 | 167 | 327.5 | No | L6139_0 |
| 276 | 797531 | 155 | 131.6 | No | L6139_1 |
| 277 | 214326 | 138 | 155.6 | No | L6148_0 |
| 278 | 632985 | 152 | 166.8 | No | L6150_3 |
| 279 | 456158 | 153 | 170.9 | No* | L6164_0 |
| 280 | 357040 | 125 | 172.5 | No* | L6164_2 |
| 281 | 146669 | 139 | 132.0 | No | L6175_0 |
| 282 | 233649 | 129 | 147.9 | No | L6175_1 |
| 283 | 359458 | 143 | 205.7 | No | L6175_4 |
| 284 | 338121 | 125 | 131.9 | No | L6175_5 |
| 285 | 591180 | 165 | 182.6 | No | L6176_1 |
| 286 | 194228 | 143 | 155.2 | No | L6176_4 |
| 287 | 541061 | 110 | 129.0 | No | L6221_0 |
| 288 | 694934 | 147 | 346.9 | No | L6221_1 |
| 289 | 451338 | 127 | 128.2 | No | L6226_1 |
| 290 | 470546 | 148 | 180.9 | No | L6226_2 |
| 291 | 471837 | 159 | 160.7 | No | L6226_4 |
| 292 | 373410 | 159 | 163.6 | No | L6227_1 |
| 293 | 333153 | 139 | 148.6 | No | L6237_0 |
| 294 | 561236 | 162 | 251.3 | No | L6238_0 |
| 295 | 319097 | 145 | 143.8 | No | L6258_1 |
| 296 | 343136 | 147 | 149.1 | No | L6265_0 |
| 297 | 491335 | 148 | 132.0 | No | L6265_2 |
| 298 | 294321 | 151 | 146.1 | No | L6274_1 |
| 299 | 515709 | 163 | 206.4 | No | L6284_0 |
| 300 | 375324 | 151 | 129.1 | No | L6298_0 |
| 301 | 183832 | 50 | 324.7 | CUT* | L6299_0 |
| 302 | 431113 | 154 | 164.2 | No | L6303_1 |
| 303 | 568249 | 166 | 170.7 | No | L6318_1 |
| 304 | 428869 | 156 | 161.6 | No | L6318_3 |
| 305 | 356277 | 152 | 164.1 | No | L6318_5 |
| 306 | 246970 | 137 | 154.1 | No | L6318_6 |
| 307 | 370605 | 130 | 161.4 | No | L6318_7 |
| 308 | 298832 | 159 | 218.2 | No | L6320_0 |
| 309 | 322660 | 148 | 125.7 | No | L6320_1 |
| 310 | 368674 | 159 | 181.1 | No | L6320_4 |
| 311 | 313595 | 142 | 137.8 | No | L6320_5 |
| 312 | 573189 | 156 | 206.1 | No | L6363_1 |
| 313 | 423988 | 143 | 137.7 | No | L6363_4 |
| 314 | 300763 | 140 | 191.1 | No | L6363_7 |
| 315 | 660322 | 162 | 179.2 | No | L6366_1 |
| 316 | 707900 | 175 | 230.6 | No | L6366_5 |
| 317 | 586157 | 152 | 196.5 | No | L6366_6 |
| 318 | 367344 | 155 | 170.1 | No | L6373_1 |
| 319 | 339814 | 136 | 145.6 | No | L6373_3 |
| 320 | 606166 | 164 | 126.0 | No | L6376_1 |
| 321 | 280977 | 151 | 156.7 | No | L6378_1 |
| 322 | 259039 | 135 | 134.7 | No | L6384_1 |
| 323 | 607912 | 146 | 176.6 | No | L6389_0 |
| 324 | 417396 | 128 | 119.9 | No | L6389_2 |
| 325 | 478614 | 146 | 184.2 | No | L6393_1 |
| 326 | 378214 | 139 | 128.2 | No | L6393_4 |
| 327 | 165538 | 145 | 150.7 | No | L6404_0 |
| 328 | 387317 | 136 | 121.9 | No | L6405_0 |
| 329 | 412241 | 151 | 181.0 | No | L6405_1 |
| 330 | 6407 | 16 | 131.9 | CUT | L6406_1 |
| 331 | 610873 | 116 | 142.8 | No* | L6406_8 |
| 332 | 362535 | 142 | 125.8 | No | L6412_1 |
| 333 | 789885 | 171 | 236.9 | No | L6420_1 |
| 334 | 666473 | 166 | 210.0 | No | L6432_0 |
| 335 | 605230 | 144 | 152.8 | No | L6432_2 |
| 336 | 252323 | 131 | 121.0 | No | L6435_1 |
| 337 | 288261 | 140 | 125.7 | No | L6439_1 |
| 338 | 411730 | 155 | 180.1 | No | L6447_1 |
| 339 | 275599 | 136 | 149.4 | No | L6448_1 |
| 340 | 593486 | 154 | 186.0 | No | L6449_0 |
| 341 | 373035 | 150 | 155.2 | No | L6450_0 |
| 342 | 652072 | 168 | 137.7 | No | L6450_1 |
| 343 | 19062 | 7 | 139.6 | CUT | L6450_2 |
| 344 | 926149 | 170 | 289.8 | No | L6450_3 |
| 345 | 623863 | 148 | 155.7 | No | L6450_4 |
| 346 | 706903 | 141 | 248.4 | No* | L6451_0 |
| 347 | 738684 | 124 | 240.6 | No* | L6451_1 |
| 348 | 257010 | 118 | 148.6 | No | L6457_2 |
| 349 | 258952 | 126 | 139.0 | No | L6460_0 |
| 350 | 357385 | 154 | 149.5 | No | L6462_0 |
| 351 | 374286 | 165 | 164.9 | No | L6462_1 |
| 352 | 1423981 | 175 | 237.9 | No | L6487_1 |
| 353 | 2779998 | 173 | 932.5 | No | L6487_3 |
| 354 | 779504 | 168 | 255.2 | No | L6494_1 |
| 355 | 442731 | 146 | 155.6 | No | L6494_4 |
| 356 | 443274 | 161 | 209.5 | No | L6499_1 |
| 357 | 363758 | 156 | 143.0 | No | L6500_1 |
| 358 | 599723 | 170 | 310.3 | No | L6500_5 |
| 359 | 513368 | 164 | 161.0 | No | L6500_6 |
| 360 | 46471 | 54 | 146.0 | CUT | L6507_2 |
| 361 | 81141 | 87 | 168.5 | CUT* | L6527_0 |
| 362 | 82673 | 87 | 120.8 | CUT* | L6527_1 |
| 363 | 38852 | 64 | 118.5 | CUT* | L6527_3 |
| 364 | 43897 | 73 | 121.5 | CUT* | L6527_4 |
| 365 | 767114 | 157 | 201.4 | No | L6528_1 |
| 366 | 464746 | 152 | 152.4 | No | L6533_1 |
| 367 | 1112383 | 172 | 305.9 | No | L6538_1 |
| 368 | 297717 | 133 | 143.3 | No | L6544_1 |
| 369 | 500076 | 162 | 232.5 | No | L6544_4 |
| 370 | 224987 | 120 | 119.9 | No | L6548_2 |
| 371 | 42430 | 56 | 124.9 | CUT | L6552_0 |
| 372 | 281786 | 117 | 140.1 | No | L6559_0 |
| 373 | 1131444 | 158 | 316.1 | No | L6559_2 |
| 374 | 438250 | 156 | 170.3 | No | L6570_1 |
| 375 | 359913 | 134 | 137.5 | No* | L6572_0 |
| 376 | 363811 | 116 | 140.9 | No | L6601_1 |
| 377 | 374476 | 127 | 175.1 | No | L6601_3 |
| 378 | 18564 | 19 | 158.9 | CUT | L6631_0 |
| 379 | 844706 | 165 | 350.0 | No | L6636_0 |
| 380 | 568973 | 126 | 252.2 | No | L6639_0 |
| 381 | 414762 | 120 | 134.3 | No* | L6639_1 |
| 382 | 458172 | 161 | 146.8 | No | L6641_1 |
| 383 | 622726 | 170 | 181.1 | No | L6641_3 |
| 384 | 442375 | 139 | 147.1 | No | L6652_4 |
| 385 | 470114 | 149 | 192.1 | No | L6652_5 |
| 386 | 394859 | 155 | 171.6 | No | L6660_0 |
| 387 | 516423 | 143 | 224.9 | No | L6667_0 |
| 388 | 245594 | 156 | 121.5 | No | L6667_1 |
| 389 | 241087 | 161 | 148.4 | No | L6667_3 |
| 390 | 134799 | 112 | 135.1 | No | L6679_0 |
| 391 | 923820 | 167 | 269.0 | No | L6717_1 |
| 392 | 462599 | 145 | 151.0 | No | L6717_3 |
| 393 | 10929 | 16 | 155.1 | CUT | L6738_0 |
| 394 | 664843 | 164 | 228.1 | No | L6779_0 |
| 395 | 856074 | 154 | 354.1 | No | L6779_2 |
| 396 | 360844 | 154 | 158.8 | No | L6780_0 |
| 397 | 541820 | 155 | 249.2 | No* | L6782_0 |
| 398 | 335149 | 122 | 122.8 | No* | L6791_1 |
| 399 | 9545 | 29 | 117.2 | CUT | L6797_2 |
| 400 | 645530 | 169 | 187.7 | No | L6825_0 |
| 401 | 519925 | 158 | 173.5 | No | L6825_1 |
| 402 | 287845 | 124 | 128.4 | No | L6848_0 |
| 403 | 393195 | 141 | 122.1 | No | L6848_2 |
| 404 | 430466 | 147 | 188.7 | No | L6854_0 |
| 405 | 288961 | 126 | 160.1 | No | L6860_2 |
| 406 | 618792 | 168 | 171.3 | No | L6883_0 |
| 407 | 659562 | 170 | 200.3 | No | L6883_1 |
| 408 | 404927 | 157 | 128.2 | No | L6913_1 |
| 409 | 420454 | 158 | 131.8 | No | L6913_3 |
| 410 | 416695 | 172 | 122.2 | No | L6913_4 |
| 411 | 276771 | 140 | 132.4 | No | L6914_2 |
| 412 | 1057876 | 167 | 194.1 | No | L6924_0 |
| 413 | 261457 | 135 | 122.7 | No | L6933_0 |
| 414 | 689505 | 152 | 172.1 | No | L6947_1 |
| 415 | 437604 | 119 | 205.5 | No | L6947_4 |
| 416 | 215595 | 97 | 128.9 | No* | L6954_0 |
| 417 | 270712 | 117 | 180.6 | No* | L6954_1 |
| 418 | 1053397 | 168 | 342.7 | No | L6955_0 |
| 419 | 172763 | 117 | 157.6 | No | L6956_1 |
| 420 | 347321 | 144 | 144.1 | No | L6961_1 |
| 421 | 536723 | 163 | 179.7 | No | L6961_2 |
| 422 | 482097 | 162 | 164.1 | No | L6968_1 |
| 423 | 139723 | 104 | 128.3 | No* | L6969_0 |
| 424 | 643558 | 148 | 209.0 | No | L6978_0 |
| 425 | 298960 | 127 | 143.7 | No | L6978_2 |
| 426 | 407125 | 147 | 196.4 | No | L6995_2 |
| 427 | 175471 | 93 | 124.9 | CUT* | L7024_0 |
| 428 | 455327 | 167 | 196.6 | No | L7029_0 |
| 429 | 483939 | 151 | 131.5 | No | L7029_1 |
| 430 | 258757 | 130 | 146.6 | No* | L7067_0 |
| 431 | 283578 | 130 | 155.4 | No* | L7120_1 |
| 432 | 611897 | 160 | 242.5 | No | L7128_2 |
| 433 | 235703 | 125 | 119.2 | No | L7135_2 |
| 434 | 349382 | 152 | 164.0 | No | L7141_1 |
| 435 | 543921 | 155 | 285.1 | No | L7174_0 |
| 436 | 454138 | 135 | 273.1 | No | L7174_1 |
| 437 | 241253 | 142 | 152.2 | No | L7174_4 |
| 438 | 461880 | 134 | 184.9 | No* | L7241_2 |
| 439 | 366982 | 148 | 119.5 | No | L7248_1 |
| 440 | 236079 | 124 | 119.9 | No | L7273_0 |
| 441 | 193793 | 132 | 157.3 | No | L7279_0 |
| 442 | 192203 | 133 | 157.5 | No | L7279_1 |
| 443 | 532614 | 134 | 137.7 | No | L7296_0 |
| 444 | 112456 | 62 | 166.5 | CUT* | L7313_1 |
| 445 | 214479 | 76 | 412.6 | CUT* | L7313_3 |
| 446 | 176615 | 68 | 181.5 | CUT* | L7313_4 |
| 447 | 920788 | 152 | 272.7 | No | L7324_1 |
| 448 | 571430 | 155 | 313.1 | No | L7572_0 |
| 449 | 336443 | 141 | 155.0 | No | L7572_1 |
| 450 | 482564 | 116 | 196.3 | No* | L7577_1 |
| 451 | 328971 | 68 | 230.8 | CUT* | L7602_1 |
| 452 | 82053 | 48 | 132.1 | CUT | L7602_15 |
| 453 | 398807 | 92 | 203.6 | CUT* | L7653_1 |
