## Supplemental Table 2 for "A target enrichment probe set for resolving the flagellate plant tree of life"

Supplemental Table 2. Samples used in the GoFlag 451 targeted enrichment experiment. A '*' by the taxon name indicated that we recovered fewer than 45 loci from the sample, and therefore, sequences from that sample were excluded from the phylogenetic analyses (Fig. 4; Supplemental Figure 2).

| **Family** | | **Taxon** | | | **Herbarium** | | **Collection number** | | **NCBI accession** | | **Sample ID** | | **Total DNA** | | **Total Library DNA** | | **Total Mapped Read Pairs** | | **Total Read Pairs** | | **% Reads Mapped** | | **Loci** | | **Conta-minant Loci** | | **Total Length Exons (bp)** |
| --- | --- | --- | --- | --- | --- | --- | --- | --- | --- | --- | --- | --- | --- | --- | --- | --- | --- | --- | --- | --- | --- | --- | --- | --- | --- | --- | --- |
|  | FERNS | |  |  | |  | |  | |  | |  | |  | |  | |  | |  | |  | |  | |  | |
|  | Aspleniaceae | | *Asplenium adiantum-nigrum* L.* | H | | Vare, H. 1766362 | | SAMN14843933 | | UFG_393201_P03_WE10 | | 3412.90 | | 1152.14 | | 5608 | | 196330 | | 0.03 | | 13 | | 0 | | 3168 | |
|  | Aspleniaceae | | *Asplenium holstii* Hieron | H | | Vare, H. 1766233 | | SAMN14888153 | | UFG_393201_P03_WF10 | | 3121.12 | | 1110.16 | | 261945 | | 1507476 | | 0.17 | | 327 | | 5 | | 66194 | |
|  | Athyriaceae | | *Diplazium hymenodes* (Mett.) Á. Löve & D. Löve | VT | | Fawcett, S. 377 | | SAMN14888154 | | UFG_393201_P03_WH09 | | 1949.39 | | 263.94 | | 117306 | | 561388 | | 0.21 | | 366 | | 8 | | 71369 | |
|  | Blechnaceae | | *Austroblechnum divergens* (Kunze) Gasper & V.A.O.Dittrich | VT | | Fawcett, S. 482 | | SAMN14888155 | | UFG_393201_P03_WA07 | | 830.65 | | 586.79 | | 433550 | | 1704712 | | 0.25 | | 419 | | 5 | | 80060 | |
|  | Blechnaceae | | *Parablechnum lineatum* (Sw.) Gasper & Salino | VT | | Fawcett, S. 477 | | SAMN14888156 | | UFG_393201_P03_WB07 | | 1709.91 | | 579.38 | | 354589 | | 1127088 | | 0.31 | | 419 | | 6 | | 79823 | |
|  | Blechnaceae | | *Blechnum polypodioides* Raddi | VT | | Fawcett, S. 442 | | SAMN14888157 | | UFG_393201_P03_WC07 | | 1630.36 | | 747.09 | | 342175 | | 1035576 | | 0.33 | | 426 | | 4 | | 80796 | |
|  | Blechnaceae | | *Lorinseria areolata* (L.) C.Presl | SEL | | Living accession: 2014-0164A | | SAMN14888158 | | UFG_393201_P03_WF06 | | 2955.08 | | 888.05 | | 36813 | | 284662 | | 0.13 | | 80 | | 0 | | 17714 | |
|  | Ophioglossaceae | | *Sceptridium multifidum* Nishida ex Tagawa | CBG | | Williams, E. 1084 | | SAMN14888159 | | UFG_393201_P03_WE06 | | 1530.48 | | 663.95 | | 292337 | | 1009460 | | 0.29 | | 402 | | 2 | | 75763 | |
|  | Cyatheaceae | | *Cyathea dombeyi* (Desv.) Lellinger | BONN, STU | | Lehnert, M. 1998 | | SAMN14888160 | | UFG_393201_P03_WH05 | | 2485.83 | | 1081.82 | | 147455 | | 629754 | | 0.23 | | 377 | | 2 | | 72648 | |
|  | Dennstaedtiaceae | | *Dennstaedtia bipinnata* (Cav.) Maxon | VT | | Fawcett, S. 470 | | SAMN14888161 | | UFG_393201_P03_WG07 | | 3192.71 | | 838.05 | | 28644 | | 291452 | | 0.10 | | 69 | | 0 | | 14115 | |
|  | Dicksoniaceae | | *Dicksonia mollis* Holttum | TAIF | | Chen, C.W. 4279 | | SAMN14888162 | | UFG_393201_P03_WA05 | | 3274.80 | | 1141.04 | | 186943 | | 863028 | | 0.22 | | 344 | | 6 | | 68288 | |
|  | Dicksoniaceae | | *Dicksonia sciurus* C. Chr. | TAIF | | Chen, C.W. 3355 | | SAMN14888163 | | UFG_393201_P03_WB05 | | 1968.90 | | 819.89 | | 78118 | | 514626 | | 0.15 | | 185 | | 4 | | 39677 | |
|  | Dicksoniaceae | | *Dicksonia stuebelii* Hieron. | BONN | | Tejedor A., s.n. | | SAMN14888164 | | UFG_393201_P03_WC05 | | 3498.11 | | 820.43 | | 124738 | | 721202 | | 0.17 | | 299 | | 6 | | 60628 | |
|  | Dicksoniaceae | | *Lophosoria christii* | BONN, STU | | Beaverhausen, A. 3812 | | SAMN14888165 | | UFG_393201_P03_WD05 | | 2323.14 | | 705.17 | | 463510 | | 1009476 | | 0.46 | | 419 | | 5 | | 79815 | |
|  | Dicksoniaceae | | *Lophosoria quadripinnata* (J.F. Gmel.) C. Chr. (#1) | SP | | Prado J.& Hirai R. 2139 | | SAMN14888166 | | UFG_393201_P03_WF05 | | 479.77 | | 929.17 | | 923505 | | 1925522 | | 0.48 | | 427 | | 6 | | 80762 | |
|  | Dicksoniaceae | | *Lophosoria quadripinnata* (J.F. Gmel.) C. Chr. (#2) | E | | 20071644 SBAR#50 | | SAMN14888167 | | UFG_393201_P03_WE05 | | 693.48 | | 771.94 | | 803407 | | 1587992 | | 0.51 | | 425 | | 5 | | 80528 | |
|  | Dicksoniaceae | | *Lophosoria revoluta* | BONN, STU | | Beaverhausen A. 3811 | | SAMN14888168 | | UFG_393201_P03_WG05 | | 1952.97 | | 881.96 | | 1653206 | | 3574690 | | 0.46 | | 428 | | 5 | | 81002 | |
|  | Dryopteridaceae | | *Ctenitis nemorosa* (Willd.) Ching* | VT | | Fawcett, S. 362 | | SAMN14888169 | | UFG_393201_P03_WE07 | | 3254.02 | | 743.11 | | 8778 | | 145580 | | 0.06 | | 26 | | 0 | | 5679 | |
|  | Dryopteridaceae | | *Dryopteris antarctica* (Baker) C. Chr. | H | | Vare, H. 1766365 | | SAMN14888170 | | UFG_393201_P03_WA10 | | 3043.53 | | 1347.09 | | 275195 | | 1031696 | | 0.27 | | 371 | | 7 | | 72487 | |
|  | Dryopteridaceae | | *Dryopteris filipaleata* J.P. Roux | H | | Fraser-Jenkins 12314 | | SAMN14888171 | | UFG_393201_P03_WB10 | | 3271.68 | | 1194.33 | | 107548 | | 443110 | | 0.24 | | 226 | | 1 | | 49482 | |
|  | Dryopteridaceae | | *Dryopteris manniana* (Hook.) C. Chr. | H | | Vare, H. 1766228 | | SAMN14888172 | | UFG_393201_P03_WC10 | | 1777.46 | | 1164.64 | | 281638 | | 944946 | | 0.30 | | 397 | | 5 | | 76790 | |
|  | Dryopteridaceae | | *Dryopteris patula* (Sw.) Underw. | NY | | Sundue, M. 1104 | | SAMN14888173 | | UFG_393201_P03_WG06 | | 76.56 | | 665.14 | | 468892 | | 1862664 | | 0.25 | | 431 | | 5 | | 81778 | |
|  | Dryopteridaceae | | *Dryopteris pentheri* (Krasser) C. Chr.* | H | | Fraser-Jenkins 12327 (H1573009 Reunion) | | SAMN14888174 | | UFG_393201_P03_WD10 | | 2475.90 | | 451.69 | | 8347 | | 96344 | | 0.09 | | 21 | | 0 | | 5151 | |
|  | Dryopteridaceae | | *Elaphoglossum aschersonii* Hieron. | NY | | Moran, R. 7645 | | SAMN14888175 | | UFG_393201_P03_WF09 | | 3059.00 | | 1182.42 | | 457724 | | 1765062 | | 0.26 | | 413 | | 7 | | 79085 | |
|  | Dryopteridaceae | | *Elaphoglossum bellermannianum* (Klotzsch) T. Moore | HUA, MO, NY | | Vasco, A. 506 | | SAMN14888176 | | UFG_393201_P03_WG09 | | 3632.39 | | 1226.71 | | 657862 | | 2560436 | | 0.26 | | 418 | | 4 | | 79777 | |
|  | Dryopteridaceae | | *Elaphoglossum engelii* (H. Karst.) Christ | HUA, NY | | Vasco, A. 794 | | SAMN14888177 | | UFG_393201_P03_WD09 | | 2667.92 | | 1103.05 | | 95877 | | 394770 | | 0.24 | | 290 | | 2 | | 58692 | |
|  | Dryopteridaceae | | *Elaphoglossum lehmannianum* Christ | MER, NY, VEN | | Vasco, A. 806 | | SAMN14888178 | | UFG_393201_P03_WH08 | | 3286.35 | | 1053.37 | | 167082 | | 685608 | | 0.24 | | 360 | | 3 | | 70811 | |
|  | Dryopteridaceae | | *Elaphoglossum novogranatense* A. Vasco | HUA, NY | | Vasco, A. 746 | | SAMN14888179 | | UFG_393201_P03_WA09 | | 3520.62 | | 1374.14 | | 399613 | | 1739426 | | 0.23 | | 416 | | 4 | | 79330 | |
|  | Dryopteridaceae | | *Elaphoglossum oreophilum* A. Vasco | HUA, MO, NY | | Vasco, A. 600 | | SAMN14888180 | | UFG_393201_P03_WB09 | | 3237.62 | | 1220.57 | | 67101 | | 335812 | | 0.20 | | 198 | | 2 | | 43039 | |
|  | Dryopteridaceae | | *Elaphoglossum ruficomum* Mickel | LPB, NY | | Labiak, P. 2896 | | SAMN14888181 | | UFG_393201_P03_WE09 | | 3102.30 | | 992.53 | | 178541 | | 691046 | | 0.26 | | 367 | | 3 | | 71182 | |
|  | Dryopteridaceae | | *Elaphoglossum yatesii* (Sodiro) Christ* | NY | | Moran, R. 6908 | | SAMN14888182 | | UFG_393201_P03_WC09 | | 3602.49 | | 1270.14 | | 11845 | | 217954 | | 0.05 | | 27 | | 1 | | 6248 | |
|  | Dryopteridaceae | | *Polystichum lachenense* (Hook.) Bedd. | NY | | 27954 | | SAMN14888183 | | UFG_393201_P03_WH06 | | 37.47 | | 406.62 | | 160427 | | 401442 | | 0.40 | | 340 | | 2 | | 66897 | |
|  | Dryopteridaceae | | *Polystichum monticola* N.C. Anthony & Schelpe | NGB | | Roux, J.P. 4715 | | SAMN14888184 | | UFG_393201_P03_WH10 | | 1494.00 | | 829.97 | | 471332 | | 943020 | | 0.50 | | 431 | | 3 | | 81744 | |
|  | Gleicheniaceae | | *Gleichenella pectinata* (Willd.) Ching | VT | | Fawcett, S. 444 | | SAMN14888185 | | UFG_393201_P03_WH07 | | 1190.21 | | 831.31 | | 422081 | | 1116308 | | 0.38 | | 347 | | 19 | | 65322 | |
|  | Polypodiaceae | | *Cochlidium serrulatum* (Sw.) L.E. Bishop | VT | | Fawcett, S. 486 | | SAMN14888186 | | UFG_393201_P03_WD07 | | 1514.09 | | 849.42 | | 431682 | | 981350 | | 0.44 | | 426 | | 5 | | 80835 | |
|  | Hymenophyllaceae | | *Vandenboschia radicans* (Sw.) Copel. | VT | | Fawcett, S. 446 | | SAMN14888187 | | UFG_393201_P03_WF08 | | 3250.09 | | 1055.04 | | 16456 | | 718132 | | 0.02 | | 57 | | 0 | | 13075 | |
|  | Lindsaeaceae | | *Sphenomeris clavata* (L.) Maxon | VT | | Fawcett, S. 450 | | SAMN14888188 | | UFG_393201_P03_WE08 | | 1658.53 | | 516.67 | | 453903 | | 1200880 | | 0.38 | | 402 | | 3 | | 75154 | |
|  | Lomariopsidaceae | | *Lomariopsis sorbifolia* (L.) Fée.* | VT | | Fawcett, S. 376 | | SAMN14888189 | | UFG_393201_P03_WC08 | | 3427.44 | | 636.74 | | 5016 | | 131356 | | 0.04 | | 11 | | 0 | | 2105 | |
|  | Marattiaceae | | *Danaea nodosa* (L.) Sm. | VT | | Fawcett, S. 475 | | SAMN14888190 | | UFG_393201_P03_WF07 | | 186.54 | | 493.74 | | 297473 | | 1516358 | | 0.20 | | 373 | | 7 | | 71850 | |
|  | Ophioglossaceae | | *Botrychium alaskense* W.H. Wagner & J.R. Grant (#1) | CBG | | Williams, E. 1082 | | SAMN14888191 | | UFG_393201_P03_WD06 | | 3699.13 | | 979.06 | | 51323 | | 557906 | | 0.09 | | 95 | | 2 | | 22254 | |
|  | Ophioglossaceae | | *Botrychium alaskense* W.H. Wagner & J.R. Grant (#2)* | CBG | | Williams, E. 1083 | | SAMN14888192 | | UFG_393201_P03_WC06 | | 3346.62 | | 525.99 | | 8657 | | 296306 | | 0.03 | | 23 | | 1 | | 5522 | |
|  | Ophioglossaceae | | *Botrychium lanceolatum* (S.G. Gmel.) Ångström* | CBG | | Williams, E. 1081 | | SAMN14888193 | | UFG_393201_P03_WB06 | | 2888.11 | | 506.37 | | 14694 | | 404640 | | 0.04 | | 29 | | 1 | | 6907 | |
|  | Ophioglossaceae | | *Botrypus virginianus* (L.) Michx. | CBG | | Williams, E. 1079 | | SAMN14888194 | | UFG_393201_P03_WA06 | | 736.20 | | 820.70 | | 503462 | | 1831710 | | 0.27 | | 412 | | 5 | | 77080 | |
|  | Pteridaceae | | *Cheilanthes hastata* Kunze | NGB | | Nicholson, G. & Roets, D. 882 | | SAMN14888195 | | UFG_393201_P03_WG10 | | 1729.43 | | 843.65 | | 748616 | | 2133598 | | 0.35 | | 434 | | 3 | | 81392 | |
|  | Pteridaceae | | *Vittaria lineata* (L.) Sm.* | VT | | Fawcett, S. 449 | | SAMN14888196 | | UFG_393201_P03_WG08 | | 3788.96 | | 1343.64 | | 30236 | | 361766 | | 0.08 | | 16 | | 0 | | 4066 | |
|  | Thelypteridaceae | | *Goniopteris abdita* (Proctor) Salino & T.E.Almeida | VT | | Fawcett, S. 457 | | SAMN14888197 | | UFG_393201_P03_WA08 | | 2131.01 | | 598.76 | | 415693 | | 1162478 | | 0.36 | | 382 | | 3 | | 74466 | |
|  | Thelypteridaceae | | *Goniopteris retroflexa* (L.) Pic. Serm | VT | | Fawcett, S. 471 | | SAMN14888198 | | UFG_393201_P03_WB08 | | 2992.49 | | 1127.19 | | 364187 | | 952390 | | 0.38 | | 431 | | 3 | | 81812 | |
|  | Thelypteridaceae | | *Macrothelypteris torresiana* (Gaudich.) Ching | VT | | Fawcett, S. 394 | | SAMN14888199 | | UFG_393201_P03_WD08 | | 3024.36 | | 762.18 | | 420359 | | 738786 | | 0.57 | | 417 | | 3 | | 79986 | |
|  | GYMNOSPERMS | |  |  | |  | |  | |  | |  | |  | |  | |  | |  | |  | |  | |  | |
|  | Araucariaceae | | *Agathis robusta* (C.Moore ex F.Muell.) F.M.Bailey | NSW | | Nagalingum, N. 43072 | | SAMN14888200 | | UFG_393201_P03_WH11 | | 3305.51 | | 1277.95 | | 58517 | | 176886 | | 0.33 | | 289 | | 0 | | 60682 | |
|  | Zamiaceae | | *Bowenia spectabilis* Hook. | NSW | | Nagalingum, N. 15-01 | | SAMN14888201 | | UFG_393201_P03_WC12 | | 3178.73 | | 965.27 | | 104301 | | 248846 | | 0.42 | | 161 | | 11 | | 35517 | |
|  | Cupressaceae | | *Callitris monticola* J.Garden | NSW | | Allen & N. Nagalingum 43077 | | SAMN14888202 | | UFG_393201_P03_WF11 | | 2966.97 | | 1009.08 | | 1019338 | | 1693380 | | 0.60 | | 435 | | 3 | | 83196 | |
|  | Zamiaceae | | *Ceratozamia kuesteriana* Regel | NSW | | Nagalingum, N. 15-60 | | SAMN14888203 | | UFG_393201_P03_WD12 | | 1975.37 | | 1064.09 | | 78566 | | 407448 | | 0.19 | | 296 | | 7 | | 60706 | |
|  | Cycadaceae | | *Cycas revoluta* Thunb. | NSW | | Nagalingum, N. 15-31 | | SAMN14888204 | | UFG_393201_P03_WG11 | | 3472.82 | | 1331.03 | | 9094 | | 100274 | | 0.09 | | 49 | | 2 | | 11525 | |
|  | Zamiaceae | | *Dioon mejiae* Standl. & L.O.Williams | NSW | | Nagalingum, N. 15-06 | | SAMN14888205 | | UFG_393201_P03_WE12 | | 2410.55 | | 1013.28 | | 810283 | | 1479716 | | 0.55 | | 378 | | 6 | | 73419 | |
|  | Ephedraceae | | *Ephedra trifurca* Torr. ex S.Watson | ALA | | Ickert-Bond, S. 753 | | SAMN14888206 | | UFG_393201_P03_WH12 | | 3572.93 | | 887.35 | | 33953 | | 142600 | | 0.24 | | 93 | | 2 | | 17920 | |
|  | Ginkgoaceae | | *Ginkgo biloba* L. | NSW | | Nagalingum, N. 15-37 | | SAMN14888207 | | UFG_393201_P03_WB11 | | 3350.78 | | 1288.51 | | 74960 | | 220228 | | 0.34 | | 321 | | 4 | | 62706 | |
|  | Gnetaceae | | *Gnetum gnemon* L. | FLAS | | Endara, L. 1647 | | SAMN14888208 | | UFG_393201_P03_WG12 | | 3616.34 | | 1264.86 | | 56311 | | 183166 | | 0.31 | | 215 | | 1 | | 44028 | |
|  | Zamiaceae | | *Macrozamia johnsonii* D.L.Jones & K.D.Hill | NSW | | Nagalingum, N. 15-20 | | SAMN14888209 | | UFG_393201_P03_WA12 | | 2605.34 | | 985.96 | | 535136 | | 1232060 | | 0.43 | | 363 | | 6 | | 70271 | |
|  | Pinaceae | | *Pinus pinaster* Aiton | NSW | | Nagalingum, N. 16-35 | | SAMN14888210 | | UFG_393201_P03_WC11 | | 2711.68 | | 1141.63 | | 171171 | | 675070 | | 0.25 | | 308 | | 1 | | 63730 | |
|  | Podocarpaceae | | *Podocarpus smithii* de Laub. | NSW | | Nagalingum, N. 16-71 | | SAMN14888211 | | UFG_393201_P03_WA11 | | 1847.08 | | 1165.82 | | 441289 | | 997798 | | 0.44 | | 436 | | 2 | | 84184 | |
|  | Sciadopityaceae | | *Sciadopitys verticillata* (Thunb.) Siebold & Zucc. | NSW | | Nagalingum, N. 16-94 | | SAMN14888212 | | UFG_393201_P03_WD11 | | 3004.85 | | 1193.03 | | 170060 | | 539192 | | 0.32 | | 374 | | 4 | | 73704 | |
|  | Taxaceae | | *Taxus baccata* L. | NSW | | Nagalingum, N. 16-96 | | SAMN14888213 | | UFG_393201_P03_WE11 | | 3226.31 | | 1426.83 | | 133181 | | 360754 | | 0.37 | | 349 | | 0 | | 70392 | |
|  | Welwitschiaceae | | *Welwitschia mirabilis* Hook.f. | FLAS | | UF Dept. Biology Greenhouse | | SAMN14888214 | | UFG_393201_P03_WF12 | | 2590.33 | | 1005.20 | | 834623 | | 1551488 | | 0.54 | | 382 | | 5 | | 71902 | |
|  | Zamiaceae | | *Zamia integrifolia* L.f. ex Aiton | NSW | | Nagalingum, N. 15-52 | | SAMN14888215 | | UFG_393201_P03_WB12 | | 3717.84 | | 1245.95 | | 68993 | | 254576 | | 0.27 | | 211 | | 5 | | 45253 | |
|  | HORNWORTS | |  |  | |  | |  | |  | |  | |  | |  | |  | |  | |  | |  | |  | |
|  | Anthocerotaceae | | *Anthoceros erectus* Kashyap | PSU | | Chantanaorrapint, S. & Suwanmala, O. 606 | | SAMN14888216 | | UFG_393201_P02_WG01 | | 841.85 | | 38.49 | | 987676 | | 1975380 | | 0.50 | | 393 | | 26 | | 74027 | |
|  | Anthocerotaceae | | *Anthoceros* tuberculatus Lehm. et Lindenb. | QFA | | Villarreal, JC. PA-16-1519 | | SAMN14888217 | | UFG_393201_P02_WC01 | | 424.73 | | 105.31 | | 673172 | | 1579606 | | 0.43 | | 380 | | 294 | | 70892 | |
|  | Leiosporocerotaceae | | *Leiosporoceros dussii* (Steph.) Hässel | QFA | | Villarreal, JC. PA-16-1508 | | SAMN14888218 | | UFG_393201_P02_WA01 | | 826.50 | | 130.01 | | 2607050 | | 3314264 | | 0.79 | | 341 | | 24 | | 64831 | |
|  | Notothyladaceae | | *Notothylas levieri* Schiffn. | PSU | | Chantanaorrapint, S. & Suwanmala, O. 607 | | SAMN14888219 | | UFG_393201_P02_WH01 | | 986.41 | | 155.95 | | 56351 | | 380986 | | 0.15 | | 207 | | 15 | | 39525 | |
|  | Anthocerotaceae | | *Phaeoceros exiguus* (Steph.) J. Haseg. | PSU | | Chantanaorrapint, S. & Suwanmala, O. 612 | | SAMN14888220 | | UFG_393201_P02_WA02 | | 1287.66 | | 33.07 | | 162664 | | 432480 | | 0.38 | | 361 | | 6 | | 69120 | |
|  | Anthocerotaceae | | *Phaeoceros himalayensis* Prosk. | PSU | | Chantanaorrapint, S. & Suwanmala, O. 651 | | SAMN14888221 | | UFG_393201_P02_WB02 | | 1398.28 | | 341.65 | | 168498 | | 373412 | | 0.45 | | 384 | | 5 | | 73431 | |
|  | Anthocerotaceae | | *Phaeoceros kashyapii* A. K. Asthana & S.C. Srivast. | PSU | | Chantanaorrapint, S. & Suwanmala, O. 718A | | SAMN14888222 | | UFG_393201_P02_WH02 | | 940.00 | | 138.25 | | 283799 | | 708310 | | 0.40 | | 391 | | 9 | | 74350 | |
|  | Anthocerotaceae | | *Phaeoceros perpusillus* Chantanaorr. | PSU | | Chantanaorrapint, S. & Suwanmala, O. 717 | | SAMN14888223 | | UFG_393201_P02_WG02 | | 857.79 | | 72.67 | | 678109 | | 1641438 | | 0.41 | | 406 | | 22 | | 76826 | |
|  | Anthocerotaceae | | *Phaeoceros* sp. 1 | PSU | | Chantanaorrapint, S. & Suwanmala, O. 716A | | SAMN14888224 | | UFG_393201_P02_WE02 | | 320.46 | | 19.33 | | 592553 | | 972772 | | 0.61 | | 396 | | 9 | | 75096 | |
|  | Anthocerotaceae | | *Phaeoceros* sp. 2 | PSU | | Chantanaorrapint, S. & Suwanmala, O. 716B | | SAMN14888225 | | UFG_393201_P02_WF02 | | 471.21 | | 9.22 | | 197300 | | 428104 | | 0.46 | | 264 | | 4 | | 50604 | |
|  | Anthocerotaceae | | *Phaeoceros* sp. 3 | PSU | | Chantanaorrapint, S. & Suwanmala, O. 604 | | SAMN14888226 | | UFG_393201_P02_WF01 | | 1574.70 | | 235.23 | | 438957 | | 1578922 | | 0.28 | | 398 | | 18 | | 75532 | |
|  | Anthocerotaceae | | *Phaeoceros* sp. 4 | FLAS | | Endara, L. 1648 | | SAMN14888227 | | UFG_393201_P02_WD01 | | 786.66 | | 132.61 | | 1277271 | | 2036552 | | 0.63 | | 406 | | 146 | | 76637 | |
|  | Dendrocerotaceae | | *Phaeomegaceros fimbriatus* (Gottsche) R. J. Duff, J. C. Villarreal, Cargill & Renzaglia | QFA | | Villarreal, JC. PA-16-1515 | | SAMN14888228 | | UFG_393201_P02_WB01 | | 1562.12 | | 135.17 | | 1432524 | | 3092390 | | 0.46 | | 411 | | 373 | | 77379 | |
|  | Phymatocerotaceae | | *Phymatoceros* sp | PSU | | Chantanaorrapint, S. & Suwanmala, O. 507 | | SAMN14888229 | | UFG_393201_P02_WE01 | | 1950.43 | | 497.77 | | 268072 | | 1066974 | | 0.25 | | 384 | | 13 | | 72913 | |
|  | LIVERWORTS | |  |  | |  | |  | |  | |  | |  | |  | |  | |  | |  | |  | |  | |
|  | Plagiochilaceae | | *Acrochila biserialis* (Lehm. & Lindenb.) Grolle | F | | Pocs 0060B | | SAMN14888230 | | UFG_393201_P02_WC08 | | 31.86 | | 177.20 | | 285863 | | 1339204 | | 0.21 | | 348 | | 12 | | 63927 | |
|  | Adelanthaceae | | *Adelanthus lindenbergianus* (Lehm.) Mitt.* | F | | Larraín, J. 37213 | | SAMN14888231 | | UFG_393201_P02_WA05 | | 59.62 | | 18.18 | | 5434 | | 157262 | | 0.03 | | 22 | | 2 | | 4117 | |
|  | Aneuraceae | | *Aneura pinguis* (L.) Dumort.* | F | | Larraín, J. 38674 | | SAMN14888232 | | UFG_393201_P02_WE07 | | 2293.70 | | 91.52 | | 542 | | 422898 | | 0.00 | | 4 | | 0 | | 718 | |
|  | Cephaloziaceae | | *Anomoclada portoricensis* (Hampe & Gottsche) Váňa | F | | Pocs 00226A | | SAMN14888233 | | UFG_393201_P02_WD08 | | 58.93 | | 98.27 | | 586044 | | 1264532 | | 0.46 | | 411 | | 11 | | 75626 | |
|  | Antheliaceae | | Anthelia juratzkana (Limpr.) Trevis.* | F | | Brinda et al. 5235 | | SAMN14888234 | | UFG_393201_P02_WB08 | | 1033.18 | | 21.17 | | 7160 | | 104136 | | 0.07 | | 39 | | 3 | | 7492 | |
|  | Balantiopsidaceae | | *Balantiopsis splendens* (Steph.) J.J. Engel & G.L. Merr. | F | | Larraín, J. 36405 | | SAMN14888235 | | UFG_393201_P02_WA03 | | 1505.89 | | 134.62 | | 640087 | | 1405682 | | 0.46 | | 407 | | 10 | | 74880 | |
|  | Lepidoziaceae | | *Bazzania bescherellei* (Steph.) Tixier | F | | Larraín, J. 36062 | | SAMN14888236 | | UFG_393201_P02_WF06 | | 35.10 | | 197.98 | | 1082916 | | 2491358 | | 0.43 | | 400 | | 135 | | 72907 | |
|  | Scapaniaceae | | *Blepharidophyllum densifolium* (Hook.) Ångström ex C. Massal. | F | | Larraín, J. 37219a | | SAMN14888237 | | UFG_393201_P02_WB05 | | 87.88 | | 259.98 | | 836291 | | 1717818 | | 0.49 | | 405 | | 12 | | 74305 | |
|  | Lejeuneaceae | | *Ceratolejeunea cornuta* (Lindenb.) Schiffner | F | | Larraín, J. 35741 | | SAMN14888238 | | UFG_393201_P02_WG05 | | 404.05 | | 13.74 | | 88052 | | 576306 | | 0.15 | | 144 | | 19 | | 25747 | |
|  | Jungermanniaceae | | *Chandonanthus hirtellus* (F. Weber) Mitt. | F | | Pocs 04121AC | | SAMN14888239 | | UFG_393201_P02_WF08 | | 4.93 | | 92.63 | | 807406 | | 1573130 | | 0.51 | | 408 | | 84 | | 75154 | |
|  | Chonecoleaceae | | *Clasmatocolea humilis* (Hook. f. & Taylor) Grolle* | F | | Larraín, J. 37644 | | SAMN14888240 | | UFG_393201_P02_WB03 | | 1063.54 | | 1.96 | | 760 | | 37132 | | 0.02 | | 4 | | 0 | | 751 | |
|  | Adelanthaceae | | *Cryptochila grandiflora* (Lindenb. & Gottsche) Grolle | F | | Larraín, J. 26449B | | SAMN14888241 | | UFG_393201_P02_WF03 | | 23.09 | | 229.77 | | 827442 | | 1860230 | | 0.44 | | 401 | | 13 | | 73998 | |
|  | Marchantiaceae | | *Dumortiera hirsuta* (Sw.) Nees | F | | Larraín, J. 38146 | | SAMN14888242 | | UFG_393201_P02_WC07 | | 1086.29 | | 640.64 | | 504742 | | 1169378 | | 0.43 | | 389 | | 10 | | 71645 | |
|  | Frullaniaceae | | *Frullania fertilis* De Not. | F | | Larraín, J. 38372 | | SAMN14888243 | | UFG_393201_P02_WD04 | | 1765.22 | | 721.60 | | 127753 | | 575830 | | 0.22 | | 305 | | 10 | | 56858 | |
|  | Lepidolaenaceae | | *Gackstroemia schwabei* (Herzog) Grolle | F | | Larraín, J. 26591 | | SAMN14888244 | | UFG_393201_P02_WG04 | | 598.32 | | 448.70 | | 563471 | | 1580652 | | 0.36 | | 393 | | 15 | | 72542 | |
|  | Acrobolbaceae | | *Goebelobryum unguiculatum* (Hook. f. & Taylor) Grolle | F | | Larraín, J. 35837 | | SAMN14888245 | | UFG_393201_P02_WE06 | | 377.57 | | 23.55 | | 311832 | | 671958 | | 0.46 | | 361 | | 13 | | 66516 | |
|  | Jungermanniaceae | | *Jamesoniella colorata* (Lehm.) Steph.* | F | | Larraín, J. 37744 | | SAMN14888246 | | UFG_393201_P02_WC04 | | 19.84 | | 69.34 | | 4606 | | 644526 | | 0.01 | | 23 | | 1 | | 4517 | |
|  | Pallavicinaceae | | *Jensenia decipiens* (Mitt.) Grolle | F | | Larraín, J. 37825 | | SAMN14888247 | | UFG_393201_P02_WG07 | | 18.46 | | 133.68 | | 547070 | | 1759926 | | 0.31 | | 361 | | 19 | | 66617 | |
|  | Trichocoleaceae | | *Leiomitra elegans* (Lehm.) Hässel de Menéndez | F | | Larraín, J. 26924 | | SAMN14888248 | | UFG_393201_P02_WB07 | | 102.36 | | 623.43 | | 244803 | | 1002712 | | 0.24 | | 374 | | 27 | | 68278 | |
|  | Lejeuneaceae | | *Lejeunea anisophylla* Mont. | F | | Larraín, J. 35316 | | SAMN14888249 | | UFG_393201_P02_WA07 | | 76.76 | | 324.36 | | 120865 | | 883266 | | 0.14 | | 151 | | 31 | | 27849 | |
|  | Pseudolepicoleaceae | | *Lepicolea rigida* (De Not.) G.A.M. Scott | F | | Larraín, J. 26518 | | SAMN14888250 | | UFG_393201_P02_WC05 | | 50.86 | | 308.49 | | 86245 | | 1085714 | | 0.08 | | 254 | | 5 | | 49214 | |
|  | Lepidoziaceae | | *Lepidozia chordulifera* Taylor | F | | Larraín, J. 26748 | | SAMN14888251 | | UFG_393201_P02_WH03 | | 16.89 | | 161.84 | | 80674 | | 616938 | | 0.13 | | 224 | | 7 | | 44119 | |
|  | Jungermanniaceae | | *Lophozia excisa* (Dicks.) Dumort. | F | | Larraín, J. 38588 | | SAMN14888252 | | UFG_393201_P02_WF05 | | 403.26 | | 913.18 | | 2273435 | | 3639930 | | 0.62 | | 417 | | 186 | | 76584 | |
|  | Aytoniaceae | | *Mannia triandra* (Scop.) Grolle | F | | Brinda et al. 6000 | | SAMN14888253 | | UFG_393201_P02_WG08 | | 1360.40 | | 105.56 | | 165874 | | 1151312 | | 0.14 | | 345 | | 25 | | 64215 | |
|  | Marchantiaceae | | *Marchantia* sp | F | | Larraín, J. 37574 | | SAMN14888254 | | UFG_393201_P02_WH04 | | 1323.69 | | 675.17 | | 1766143 | | 2379170 | | 0.74 | | 407 | | 10 | | 74826 | |
|  | Lejeuneaceae | | *Mastigolejeunea auriculata* (Wilson & Hook.) Schiffner | F | | Larraín, J. 35461 | | SAMN14888255 | | UFG_393201_P02_WD06 | | 131.01 | | 177.71 | | 370846 | | 951296 | | 0.39 | | 248 | | 27 | | 45241 | |
|  | Jungermanniaceae | | *Mesoptychia badensis* (Gottsche ex Rabenh.) L.Soederstr. & Vana* | F | | Brinda 6464 | | SAMN14888256 | | UFG_393201_P02_WE08 | | 2264.37 | | 16.17 | | 17234 | | 170342 | | 0.10 | | 3 | | 101 | | 605 | |
|  | Metzgeriaceae | | *Metzgeria decrescens* Steph* | F | | Larraín, J. 27039B | | SAMN14888257 | | UFG_393201_P02_WG06 | | 51.94 | | 11.69 | | 3420 | | 118766 | | 0.03 | | 13 | | 1 | | 2363 | |
|  | Adelanthaceae | | *Odontoschisma navicularis* (Steph.) Grolle | F | | Larraín, J. 35892 | | SAMN14888258 | | UFG_393201_P02_WD07 | | 1.18 | | 35.20 | | 335021 | | 891094 | | 0.38 | | 398 | | 5 | | 73537 | |
|  | Plagiochilaceae | | *Pedinophyllopsis abdita* (Sull.) R.M. Schust. & Inoue | F | | Larraín, J. 37607 | | SAMN14888259 | | UFG_393201_P02_WE04 | | 665.89 | | 85.34 | | 294004 | | 1155834 | | 0.25 | | 373 | | 15 | | 67979 | |
|  | Aytoniaceae | | *Plagiochasma cordatum* Lehm. & Lindenb. | PSU | | Chantanaorrapint, S. & Suwanmala, O. 678 | | SAMN14888260 | | UFG_393201_P02_WD02 | | 1945.06 | | 341.65 | | 83211 | | 421472 | | 0.20 | | 254 | | 10 | | 49066 | |
|  | Plagiochilaceae | | *Plagiochila exigua* (Taylor) Taylor | F | | Pocs 05026B | | SAMN14888261 | | UFG_393201_P02_WA06 | | 42.59 | | 309.52 | | 163326 | | 954240 | | 0.17 | | 295 | | 103 | | 55242 | |
|  | Porellaceae | | *Porella recurva* (Taylor) Kuhnem. | F | | Larraín, J. 26650 | | SAMN14888262 | | UFG_393201_P02_WB04 | | 1149.80 | | 608.17 | | 164642 | | 406308 | | 0.41 | | 364 | | 4 | | 68521 | |
|  | Ptilidiaceae | | *Ptilidium ciliare* # 1 (L.) Hampe | F | | Larraín, J. 33007 | | SAMN14888263 | | UFG_393201_P02_WD05 | | 1968.44 | | 675.67 | | 76261 | | 382086 | | 0.20 | | 220 | | 8 | | 43504 | |
|  | Ptilidiaceae | | *Ptilidium ciliare* # 2 (L.) Hampe | FLAS | | Moss Dimensions Team 153 | | SAMN14888264 | | UFG_393201_P02_WB11 | | 136.43 | | 384.74 | | 971927 | | 1699332 | | 0.57 | | 405 | | 13 | | 74072 | |
|  | Radulaceae | | *Radula novocaledoniensis* K. Yamada | F | | Larraín, J. 35812 | | SAMN14888265 | | UFG_393201_P02_WB06 | | 58.44 | | 388.15 | | 186622 | | 1144476 | | 0.16 | | 334 | | 12 | | 61897 | |
|  | Aytoniaceae | | *Reboulia hemisphaerica* (L.) Raddi | F | | Larraín, J. 36056 | | SAMN14888266 | | UFG_393201_P02_WC06 | | 709.30 | | 550.63 | | 464860 | | 1405798 | | 0.33 | | 381 | | 14 | | 69718 | |
|  | Aneuraceae | | *Riccardia spegazziniana* A. Massal. | F | | Larraín, J. 36876 | | SAMN14888267 | | UFG_393201_P02_WF04 | | 800.98 | | 24.02 | | 5038 | | 68976 | | 0.07 | | 26 | | 3 | | 5091 | |
|  | Ricciaceae | | *Riccia atromarginata* Levier | F | | Pocs 03024D | | SAMN14888268 | | UFG_393201_P02_WH06 | | 194.32 | | 447.38 | | 29978 | | 723718 | | 0.04 | | 146 | | 7 | | 28223 | |
|  | Adelanthaceae | | *Syzygiella jacquinotii* (Mont.) Hentschel, K. Feldberg, Váňa & Heinrichs | F | | Larraín, J. 37233 | | SAMN14888269 | | UFG_393201_P02_WD03 | | 1091.02 | | 354.36 | | 379862 | | 1181604 | | 0.32 | | 395 | | 9 | | 73222 | |
|  | Geocalycaceae | | *Saccogynidium vasculosum* (Hook. f. & Taylor) Grolle | F | | Larraín, J. 34239 | | SAMN14888270 | | UFG_393201_P02_WH07 | | 66.12 | | 40.24 | | 173513 | | 446496 | | 0.39 | | 337 | | 7 | | 62412 | |
|  | Schistochilaceae | | *Schistochila acuminata* Steph. | F | | Larraín, J. 37847 | | SAMN14888271 | | UFG_393201_P02_WE05 | | 812.29 | | 354.36 | | 580686 | | 1635594 | | 0.36 | | 406 | | 6 | | 74294 | |
|  | Targioniaceae | | *Targionia hypophylla* L. (#1) | F | | Pocs 04029J | | SAMN14888272 | | UFG_393201_P02_WC02 | | 63.86 | | 48.73 | | 209233 | | 506806 | | 0.41 | | 338 | | 31 | | 62965 | |
|  | Targioniaceae | | *Targionia hypophylla* L. (#2) | PSU | | Chantanaorrapint, S. & Suwanmala, O. 668 | | SAMN14888273 | | UFG_393201_P02_WA08 | | 2615.04 | | 275.77 | | 498664 | | 1056906 | | 0.47 | | 394 | | 9 | | 72329 | |
|  | Herbertaceae | | *Triandrophyllum* sp | F | | Larraín, J. 37419 | | SAMN14888274 | | UFG_393201_P02_WA04 | | 67.30 | | 234.93 | | 94031 | | 608080 | | 0.15 | | 275 | | 6 | | 53390 | |
|  | Acrobolbaceae | | *Acrobolbus urvilleanus* (Mont.) Trevis | F | | Larraín, J. 27111 | | SAMN14888275 | | UFG_393201_P02_WH05 | | 191.86 | | 440.85 | | 127112 | | 640582 | | 0.20 | | 358 | | 7 | | 66009 | |
|  | LYCOPHYTES | |  |  | |  | |  | |  | |  | |  | |  | |  | |  | |  | |  | |  | |
|  | Lycopodiaceae | | *Austrolycopodium magellanicum* (P. Beauv.) Holub | HUA, VT | | Testo, W. 966 | | SAMN14888276 | | UFG_393201_P03_WF03 | | 959.74 | | 804.10 | | 745662 | | 1319374 | | 0.57 | | 418 | | 5 | | 78051 | |
|  | Lycopodiaceae | | *Dendrolycopodium dendroideum* (Michx.) A. Haines | VT | | Testo, W. 405 | | SAMN14888277 | | UFG_393201_P03_WG03 | | 263.94 | | 1236.30 | | 1042660 | | 2380654 | | 0.44 | | 420 | | 4 | | 78243 | |
|  | Lycopodiaceae | | *Diphasiastrum digitatum* (Dill.) Holub | VT | | Testo, W. 404 | | SAMN14888278 | | UFG_393201_P03_WH03 | | 1267.22 | | 869.68 | | 420511 | | 742458 | | 0.57 | | 412 | | 5 | | 76929 | |
|  | Lycopodiaceae | | *Diphasium scariosum* (Forst.) Rothm. | VT | | Sundue, M. 3648 | | SAMN14888279 | | UFG_393201_P03_WA04 | | 2621.04 | | 865.05 | | 296178 | | 618202 | | 0.48 | | 362 | | 6 | | 69380 | |
|  | Lycopodiaceae | | *Huperzia miyoshiana* (Makino) Ching | DUKE | | Rothfels, C. 4483 | | SAMN14888280 | | UFG_393201_P03_WE04 | | 1389.38 | | 492.18 | | 796076 | | 1660358 | | 0.48 | | 425 | | 3 | | 79408 | |
|  | Isoetaceae | | *Isoetes echinospora* Durieu | S | | Hammarsjö nb03 1913 | | SAMN14888281 | | UFG_393201_P03_WD03 | | 323.02 | | 693.16 | | 546206 | | 981034 | | 0.56 | | 413 | | 5 | | 76907 | |
|  | Isoetaceae | | *Isoetes lacustris* L. | S | | Haraldsson, et al # nb06 1362 | | SAMN14888282 | | UFG_393201_P03_WE03 | | 1444.35 | | 88.64 | | 524633 | | 813542 | | 0.64 | | 406 | | 3 | | 76093 | |
|  | Lycopodiaceae | | *Lycopodium clavatum* L. | MEXU, VT | | Matos 2462 | | SAMN14888283 | | UFG_393201_P03_WH04 | | 2419.67 | | 927.93 | | 67754 | | 238872 | | 0.28 | | 191 | | 1 | | 41340 | |
|  | Lycopodiaceae | | *Lycopodium clavatum* subsp. *contiguum* (Klotzsch) B. Øllg. | CR, VT | | Testo, W. 695 | | SAMN14888284 | | UFG_393201_P03_WC04 | | 124.51 | | 598.13 | | 745957 | | 1577866 | | 0.47 | | 422 | | 8 | | 78731 | |
|  | Lycopodiaceae | | *Lycopodium lagopus* (Laestadius ex C. Hartman) G. Zinserling ex Kuzeneva-Prochorova | VT | | Testo, W. 597 | | SAMN14888285 | | UFG_393201_P03_WD04 | | 2273.38 | | 933.64 | | 295808 | | 547212 | | 0.54 | | 394 | | 6 | | 74647 | |
|  | Lycopodiaceae | | *Palhinhaea tomentosa* (Alderw.) Holub | VT | | Sundue, M. 3718 | | SAMN14888286 | | UFG_393201_P03_WG04 | | 1116.54 | | 963.38 | | 897491 | | 2588352 | | 0.35 | | 382 | | 16 | | 70235 | |
|  | Lycopodiaceae | | *Phlegmariurus lauterbachii* (E.Pritz. ex K.Schum. & Lauterb.) A.R.Field & Bostock | Z | | Karger 2560 | | SAMN14888287 | | UFG_393201_P03_WF04 | | 2320.49 | | 1093.94 | | 224406 | | 914080 | | 0.25 | | 364 | | 2 | | 69496 | |
|  | Lycopodiaceae | | *Pseudodiphasium volubile* (G. Forst.) Holub | VT | | Sundue, M. 3614 | | SAMN14888288 | | UFG_393201_P03_WB04 | | 914.25 | | 754.80 | | 954597 | | 1476690 | | 0.65 | | 423 | | 4 | | 79184 | |
|  | Selaginellaceae | | *Selaginella estrellensis* Hieron. | SEL | | Living accession: 1986-0492A | | SAMN14888289 | | UFG_393201_P03_WC03 | | 554.61 | | 978.25 | | 2617590 | | 3956478 | | 0.66 | | 396 | | 9 | | 73419 | |
|  | Selaginellaceae | | *Selaginella plana* (Desv. ex Poir.) Hieron. | VT | | Fawcett, S. 481 | | SAMN14888290 | | UFG_393201_P03_WA03 | | 1750.79 | | 534.31 | | 961168 | | 1699118 | | 0.57 | | 325 | | 23 | | 60615 | |
|  | Selaginellaceae | | *Selaginella selaginoides* (L.) P. Beauv. ex Mart. & Schrank | S | | Haraldsson, Kerstin & Asplund, Gunvor # nb06 1335 | | SAMN14888291 | | UFG_393201_P03_WB03 | | 502.52 | | 386.58 | | 4212119 | | 5559842 | | 0.76 | | 324 | | 25 | | 60381 | |
|  | MOSSES | |  |  | |  | |  | |  | |  | |  | |  | |  | |  | |  | |  | |  | |
|  | Polytrichaceae | | *Atrichum angustatum* (Brid.) Bruch & Schimp. | Culture collection Rensing Lab | | CCRL69 (NC991): Jesson, L. | | SAMN14888292 | | UFG_393201_P03_WB01 | | 1866.14 | | 705.01 | | 1342788 | | 2018568 | | 0.67 | | 398 | | 2 | | 72792 | |
|  | Polytrichaceae | | *Atrichum undulatum* (Hedw.) P. Beauv. | Culture collection Rensing Lab | | CCRL70: Beike A.K. | | SAMN14888293 | | UFG_393201_P03_WC01 | | 2421.98 | | 833.74 | | 2583119 | | 3315080 | | 0.78 | | 405 | | 5 | | 73574 | |
|  | Aulacomniaceae | | *Aulacomnium acuminatum* (Lindb. & Arnell) Kindb. | FLAS | | Moss Dimensions Team 125 | | SAMN14888294 | | UFG_393201_P02_WA09 | | 399.33 | | 1007.00 | | 3687765 | | 4418854 | | 0.83 | | 415 | | 9 | | 76018 | |
|  | Aulacomniaceae | | *Aulacomnium androgynum* (Hedw.) Schwägr. | Culture collection Rensing Lab | | CCRL771: Beike A.K. | | SAMN14888295 | | UFG_393201_P03_WD01 | | 396.96 | | 1176.81 | | 5986206 | | 6725502 | | 0.89 | | 417 | | 2 | | 76178 | |
|  | Aulacomniaceae | | *Aulacomnium palustre* (Hedw.) Schwägr. | FLAS | | Moss Dimensions Team 363 | | SAMN14888296 | | UFG_393201_P02_WH08 | | 231.64 | | 853.79 | | 5753664 | | 6550852 | | 0.88 | | 417 | | 3 | | 76525 | |
|  | Aulacomniaceae | | *Aulacomnium turgidum* (Wahlenb.) Schwägr. | FLAS | | Moss Dimensions Team 126 | | SAMN14888297 | | UFG_393201_P02_WB09 | | 472.58 | | 921.54 | | 5638969 | | 6612900 | | 0.85 | | 418 | | 3 | | 76458 | |
|  | Ditrichaceae | | *Ceratodon purpureus* (Hedw.) Brid. | NYS | | Miller, N.G.18,934 | | SAMN14888298 | | UFG_393201_P03_WG02 | | 1798.47 | | 403.72 | | 1622780 | | 2438000 | | 0.67 | | 416 | | 2 | | 76486 | |
|  | Dicranaceae | | *Dicranella heteromalla* (Hedw.) Schimp. | Culture collection Rensing Lab | | CCRL82: Beike A.K. | | SAMN14888299 | | UFG_393201_P03_WE01 | | 1050.15 | | 163.81 | | 939520 | | 1395520 | | 0.67 | | 417 | | 3 | | 76267 | |
|  | Dicranaceae | | *Dicranum acutifolium* (Lindb. & Arnell) C.E.O. Jensen | FLAS | | Moss Dimensions Team 394 | | SAMN14888300 | | UFG_393201_P02_WF10 | | 1358.79 | | 909.83 | | 6687466 | | 8758348 | | 0.76 | | 416 | | 10 | | 76480 | |
|  | Dicranaceae | | *Dicranum elongatum* Schleich. ex Schwägr. | FLAS | | Moss Dimensions Team 395 | | SAMN14888301 | | UFG_393201_P02_WH10 | | 559.23 | | 603.47 | | 2041066 | | 2793428 | | 0.73 | | 414 | | 22 | | 76222 | |
|  | Dicranaceae | | *Dicranum fragilifolium* Lindb. | FLAS | | Moss Dimensions Team 396 | | SAMN14888302 | | UFG_393201_P02_WD10 | | 910.44 | | 579.45 | | 2287613 | | 3205920 | | 0.71 | | 414 | | 9 | | 76254 | |
|  | Dicranaceae | | *Dicranum montanum* Hedw. | FLAS | | Moss Dimensions Team 397 | | SAMN14888303 | | UFG_393201_P02_WA11 | | 1833.46 | | 834.93 | | 2394528 | | 3300318 | | 0.73 | | 415 | | 5 | | 76517 | |
|  | Dicranaceae | | *Dicranum polysetum* Sw. | FLAS | | Moss Dimensions Team 398 | | SAMN14888304 | | UFG_393201_P02_WC10 | | 1907.24 | | 704.54 | | 141852 | | 517710 | | 0.27 | | 283 | | 2 | | 55751 | |
|  | Dicranaceae | | *Dicranum scoparium* Hedw. | FLAS | | Moss Dimensions Team 399 | | SAMN14888305 | | UFG_393201_P02_WG10 | | 1755.41 | | 544.09 | | 787325 | | 1983888 | | 0.40 | | 409 | | 6 | | 75524 | |
|  | Dicranaceae | | *Dicranum undulatum* Schrad. ex Brid. | FLAS | | Moss Dimensions Team 400 | | SAMN14888306 | | UFG_393201_P02_WE10 | | 1483.37 | | 719.10 | | 4084744 | | 5362868 | | 0.76 | | 416 | | 10 | | 76364 | |
|  | Encalyptaceae | | *Encalypta vulgaris* Hedw. | Culture collection Rensing Lab | | CCRL84: Frahm J.-P. | | SAMN14888307 | | UFG_393201_P03_WF01 | | 48.50 | | 342.88 | | 6587108 | | 7641178 | | 0.86 | | 418 | | 2 | | 76956 | |
|  | Funariaceae | | *Entosthodon smithhurstii* (Broth. & Geh.) Paris | Culture collection Rensing Lab | | CCRL91: Goffinet B. | | SAMN14888308 | | UFG_393201_P03_WG01 | | 729.39 | | 413.78 | | 6275109 | | 7438474 | | 0.84 | | 421 | | 2 | | 78879 | |
|  | Fissidentaceae | | *Fissidens bryoides* Hedw. | Culture collection Rensing Lab | | CCRL92: Beike A.K. | | SAMN14888309 | | UFG_393201_P03_WH01 | | 817.95 | | 194.23 | | 3140664 | | 3581912 | | 0.88 | | 414 | | 5 | | 75567 | |
|  | Funariaceae | | *Funaria hygrometrica* Hedw. | Culture collection Rensing Lab | | CCRL6: Frahm J.-P. | | SAMN14888310 | | UFG_393201_P03_WA01 | | 259.90 | | 626.77 | | 2240300 | | 2780802 | | 0.81 | | 420 | | 2 | | 78044 | |
|  | Hylocomiaceae | | *Hylocomium splendens* (Hedw.) Schimp. | FLAS | | Moss Dimensions Team 14 | | SAMN14888311 | | UFG_393201_P02_WG09 | | 325.28 | | 1143.49 | | 435445 | | 686456 | | 0.63 | | 417 | | 3 | | 76634 | |
|  | Hypnaceae | | *Hypnum lindbergii* Mitt. | FLAS | | Moss Dimensions Team 401 | | SAMN14888312 | | UFG_393201_P02_WF11 | | 1768.80 | | 710.99 | | 3292582 | | 4423302 | | 0.74 | | 420 | | 7 | | 77286 | |
|  | Leucobryaceae | | *Leucobryum glaucum* (Hedw.) Ångstr. | Culture collection Rensing Lab | | CCRL94: Beike A.K. | | SAMN14888313 | | UFG_393201_P03_WA02 | | 51.25 | | 455.05 | | 3731367 | | 4484574 | | 0.83 | | 421 | | 4 | | 77772 | |
|  | Grimmiaceae | | *Racomitrium canescens* (Hedw.) Bednarek-Ochyra & Ochyra |  | | S. McDaniel (pending) MossDim2016. 12.20.Racspp | | SAMN14888314 | | UFG_393201_P02_WH09 | | 301.26 | | 756.52 | | 505609 | | 1148668 | | 0.44 | | 412 | | 0 | | 76013 | |
|  | Funariaceae | | *Physcomitrella patens* (Hedw.) Bruch & Schimp. | Culture collection Rensing Lab | | CCRL16: Gransden | | SAMN14888315 | | UFG_393201_P03_WH02 | | 877.53 | | 395.45 | | 2588376 | | 3047860 | | 0.85 | | 420 | | 4 | | 78551 | |
|  | Mniaceae | | *Plagiomnium undulatum* (Hedw.) T.J. Kop. | Culture collection Rensing Lab | | CCRL105: Beike A.K. | | SAMN14888316 | | UFG_393201_P03_WB02 | | 73.90 | | 464.42 | | 3866506 | | 4929996 | | 0.78 | | 418 | | 4 | | 76215 | |
|  | Hylocomiaceae | | *Pleurozium schreberi* (Willd. ex Brid.) Mitt. | FLAS | | Moss Dimensions Team 71 | | SAMN14888317 | | UFG_393201_P02_WB10 | | 563.66 | | 881.06 | | 822851 | | 1391112 | | 0.59 | | 417 | | 4 | | 76398 | |
|  | Polytrichaceae | | *Polytrichum commune* Hedw. | FLAS | | Moss Dimensions Team 384 | | SAMN14888318 | | UFG_393201_P02_WD09 | | 478.59 | | 496.27 | | 867818 | | 1391264 | | 0.62 | | 405 | | 4 | | 73114 | |
|  | Polytrichaceae | | Polytrichum juniperinum Hedw. | FLAS | | Moss Dimensions Team 389 | | SAMN14888319 | | UFG_393201_P02_WC09 | | 488.44 | | 766.78 | | 1112429 | | 1955284 | | 0.57 | | 407 | | 7 | | 73883 | |
|  | Polytrichaceae | | *Polytrichum strictum* Menzies ex Brid. | FLAS | | Moss Dimensions Team 106 | | SAMN14888320 | | UFG_393201_P02_WE09 | | 504.78 | | 612.27 | | 2460723 | | 3177384 | | 0.77 | | 408 | | 9 | | 73821 | |
|  | Hypnaceae | | *Ptilium crista-castrensis* (Hedw.) De Not. | FLAS | | Moss Dimensions Team 42 | | SAMN14888321 | | UFG_393201_P02_WC11 | | 1263.30 | | 881.61 | | 3479835 | | 4312976 | | 0.81 | | 418 | | 2 | | 76830 | |
|  | Grimmiaceae | | *Racomitrium lanuginosum* (Hedw.) Brid. |  | | S. McDaniel (pending) | | SAMN14888322 | | UFG_393201_P02_WA10 | | 1132.36 | | 631.54 | | 1164994 | | 1773390 | | 0.66 | | 421 | | 3 | | 77410 | |
|  | Brachytheciaceae | | Rhynchostegium murale (Hedw.) Schimp. | Culture collection Rensing Lab | | CCRL109: Beike A.K. | | SAMN14888323 | | UFG_393201_P03_WC02 | | 1199.21 | | 502.56 | | 3711502 | | 4126782 | | 0.90 | | 418 | | 4 | | 76729 | |
|  | Hylocomiaceae | | *Rhytidiadelphus triquetrus* (Hedw.) Warnst. |  | | S. McDaniel (pending) | | SAMN14888324 | | UFG_393201_P02_WE11 | | 912.63 | | 818.42 | | 3285077 | | 4740388 | | 0.69 | | 419 | | 11 | | 77057 | |
|  | Hylocomiaceae | | *Rhytidium rugosum* (Ehrh. ex Hedw.) Kindb. | FLAS | | Moss Dimensions Team 164 | | SAMN14888325 | | UFG_393201_P02_WD11 | | 1048.07 | | 951.56 | | 2783364 | | 3726078 | | 0.75 | | 417 | | 10 | | 76820 | |
|  | Sphagnaceae | | *Sphagnum alaskense* R.E. Andrus & Janssens | FLAS | | Moss Dimensions Team 402 | | SAMN14888326 | | UFG_393201_P02_WG11 | | 1215.03 | | 942.25 | | 2580712 | | 3139982 | | 0.82 | | 409 | | 8 | | 74232 | |
|  | Sphagnaceae | | *Sphagnum angustifolium* (Warnst.) C.E.O. Jensen | FLAS | | Moss Dimensions Team 403 | | SAMN14888327 | | UFG_393201_P02_WH12 | | 1487.07 | | 280.37 | | 687925 | | 1065912 | | 0.65 | | 406 | | 3 | | 73391 | |
|  | Sphagnaceae | | *Sphagnum arcticum* Flatberg & Frisvoll | FLAS | | Moss Dimensions Team 404 | | SAMN14888328 | | UFG_393201_P02_WD12 | | 1692.13 | | 687.53 | | 1407820 | | 1993264 | | 0.71 | | 407 | | 11 | | 73394 | |
|  | Sphagnaceae | | *Sphagnum compactum* Lam. & DC. | FLAS | | Moss Dimensions Team 405 | | SAMN14888329 | | UFG_393201_P02_WA12 | | 2305.02 | | 762.38 | | 1942253 | | 2357978 | | 0.82 | | 410 | | 9 | | 74171 | |
|  | Sphagnaceae | | *Sphagnum fimbriatum* Wilson | FLAS | | Moss Dimensions Team 406 | | SAMN14888330 | | UFG_393201_P02_WC12 | | 811.37 | | 792.65 | | 2368258 | | 3061904 | | 0.77 | | 408 | | 9 | | 73539 | |
|  | Sphagnaceae | | *Sphagnum fuscum* (Schimp.) H. Klinggr. | FLAS | | Moss Dimensions Team 407 | | SAMN14888331 | | UFG_393201_P02_WE12 | | 1497.46 | | 672.72 | | 1019764 | | 1297964 | | 0.79 | | 405 | | 7 | | 73042 | |
|  | Sphagnaceae | | *Sphagnum girgensohnii* Russow | FLAS | | Moss Dimensions Team 408 | | SAMN14888332 | | UFG_393201_P02_WG12 | | 1965.67 | | 624.58 | | 621644 | | 989484 | | 0.63 | | 406 | | 7 | | 73302 | |
|  | Sphagnaceae | | *Sphagnum magellanicum* Brid. | FLAS | | Moss Dimensions Team 409 | | SAMN14888333 | | UFG_393201_P02_WH11 | | 1518.82 | | 707.89 | | 294837 | | 440046 | | 0.67 | | 393 | | 4 | | 71812 | |
|  | Sphagnaceae | | *Sphagnum russowii* Warnst. | FLAS | | Moss Dimensions Team 410 | | SAMN14888334 | | UFG_393201_P02_WF12 | | 1839.23 | | 736.51 | | 2402028 | | 3441872 | | 0.70 | | 408 | | 11 | | 73544 | |
|  | Sphagnaceae | | *Sphagnum squarrosum* Crome | FLAS | | Moss Dimensions Team 411 | | SAMN14888335 | | UFG_393201_P02_WB12 | | 2320.72 | | 778.79 | | 1519007 | | 2116438 | | 0.72 | | 408 | | 8 | | 74296 | |
|  | Takakiaceae | | *Takakia lepizioides* S. Hatt. & Inoue | Culture collection Rensing Lab | | CCRL113: Li, X. & Hu, R. | | SAMN14888336 | | UFG_393201_P03_WD02 | | 1073.13 | | 469.11 | | 2045612 | | 2783936 | | 0.73 | | 384 | | 16 | | 70853 | |
|  | Thuidiaceae | | *Thuidium tamariscinum* (Hedw.) Schimp. | Culture collection Rensing Lab | | CCRL114: Beike A.K. | | SAMN14888337 | | UFG_393201_P03_WE02 | | 1843.74 | | 1020.12 | | 4321099 | | 4846942 | | 0.89 | | 416 | | 3 | | 76752 | |
|  | Brachytheciaceae | | *Tomentypnum nitens* (Hedw.) Loeske | FLAS | | Moss Dimensions Team 169 | | SAMN14888338 | | UFG_393201_P02_WF09 | | 157.10 | | 992.09 | | 528434 | | 990910 | | 0.53 | | 410 | | 4 | | 75567 | |
|  | Pottiaceae | | *Tortula truncata* (Hedw.) Mitt. | Culture collection Rensing Lab | | CCRL115: Frahm J.-P. | | SAMN14888339 | | UFG_393201_P03_WF02 | | 1160.19 | | 142.26 | | 3639771 | | 4387540 | | 0.83 | | 418 | | 3 | | 75165 | |
