## Supplemental Table 3 for "A target enrichment probe set for resolving the flagellate plant tree of life"

**Supplemental Table 3.** Comparison of the number of loci covered by the GoFlag 451 probe set that were recovered by targeted enrichment ("GoFlag Loci") and those recovered by the 1KP transcriptome sequencing (One Thousand Plant Transcriptomes Initiative, 2019).

| **Taxon** | **GoFlag ID** | **1KP Code** | **GoFlag Loci** | **1KP Loci** |
| --- | --- | --- | --- | --- |
| *Agathis_robusta* | UFG_393201_P03_WH11 | MIXZ | 289 | 331 |
| *Atrichum_angustatum* | UFG_393201_P03_WB01 | ZTHV | 398 | 360 |
| *Botrypus_virginianus* | UFG_393201_P03_WA06 | BEGM | 412 | 284 |
| *Ceratodon_purpureus* | UFG_393201_P03_WG02 | FFPD | 416 | 369 |
| *Danaea_nodosa* | UFG_393201_P03_WF07 | DFHO | 373 | 396 |
| *Dicranum_scoparium* | UFG_393201_P02_WG10 | NGTD | 409 | 360 |
| *Diphasiastrum_digitatum* | UFG_393201_P03_WH03 | WAFT | 412 | 357 |
| *Ginkgo_biloba* | UFG_393201_P03_WB11 | SGTW | 321 | 252 |
| *Leiosporoceros_dussii* | UFG_393201_P02_WA01 | ANON | 341 | 283 |
| *Leucobrym_glaucum* | UFG_393201_P03_WA02 | RGKI | 421 | 350 |
| *Takakia_lepizioides* | UFG_393201_P02_WD09 | SZYG | 384 | 301 |
| *Sciadopitys_verticillata* | UFG_393201_P03_WD11 | YFZK | 374 | 366 |
| *Selaginella_selaginelloides* | UFG_393201_P03_WB03 | KUXM | 324 | 336 |
| *Takakia_lepizioides* | UFG_393201_P03_WD02 | SKQD | 384 | 382 |
| *Taxus_baccata* | UFG_393201_P03_WE11 | WWSS | 349 | 344 |
| *Vittaria_lineata* | UFG_393201_P03_WG08 | SKYV | 16 | 340 |
| *Welwitschia_mirabilis* | UFG_393201_P03_WF12 | TOXE | 382 | 381 |
