## Supplementary figures and images for "A target enrichment probe set for resolving the flagellate plant tree of life"

### Supplemental Figure 1

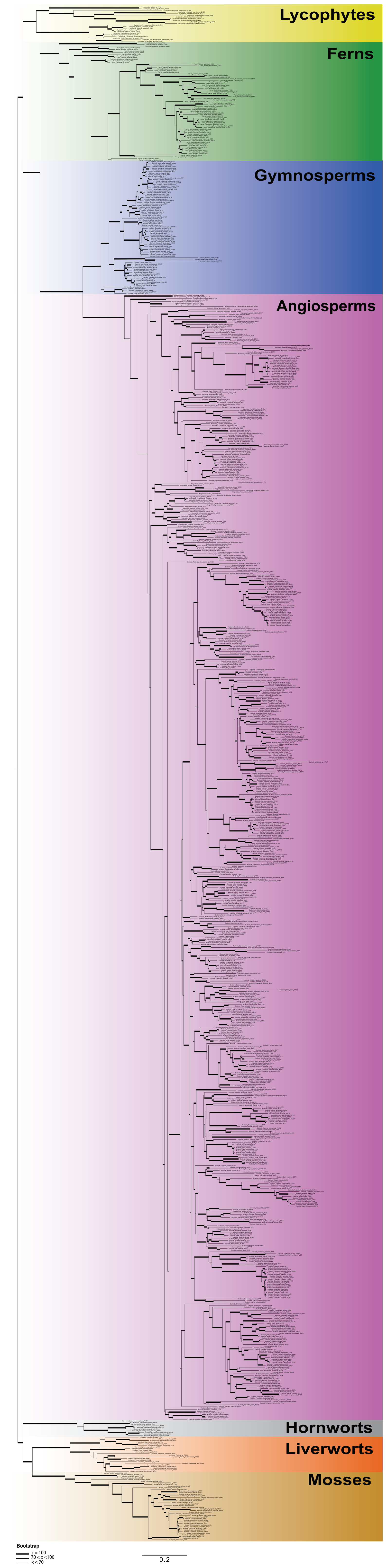

### Supplemental Figure 2

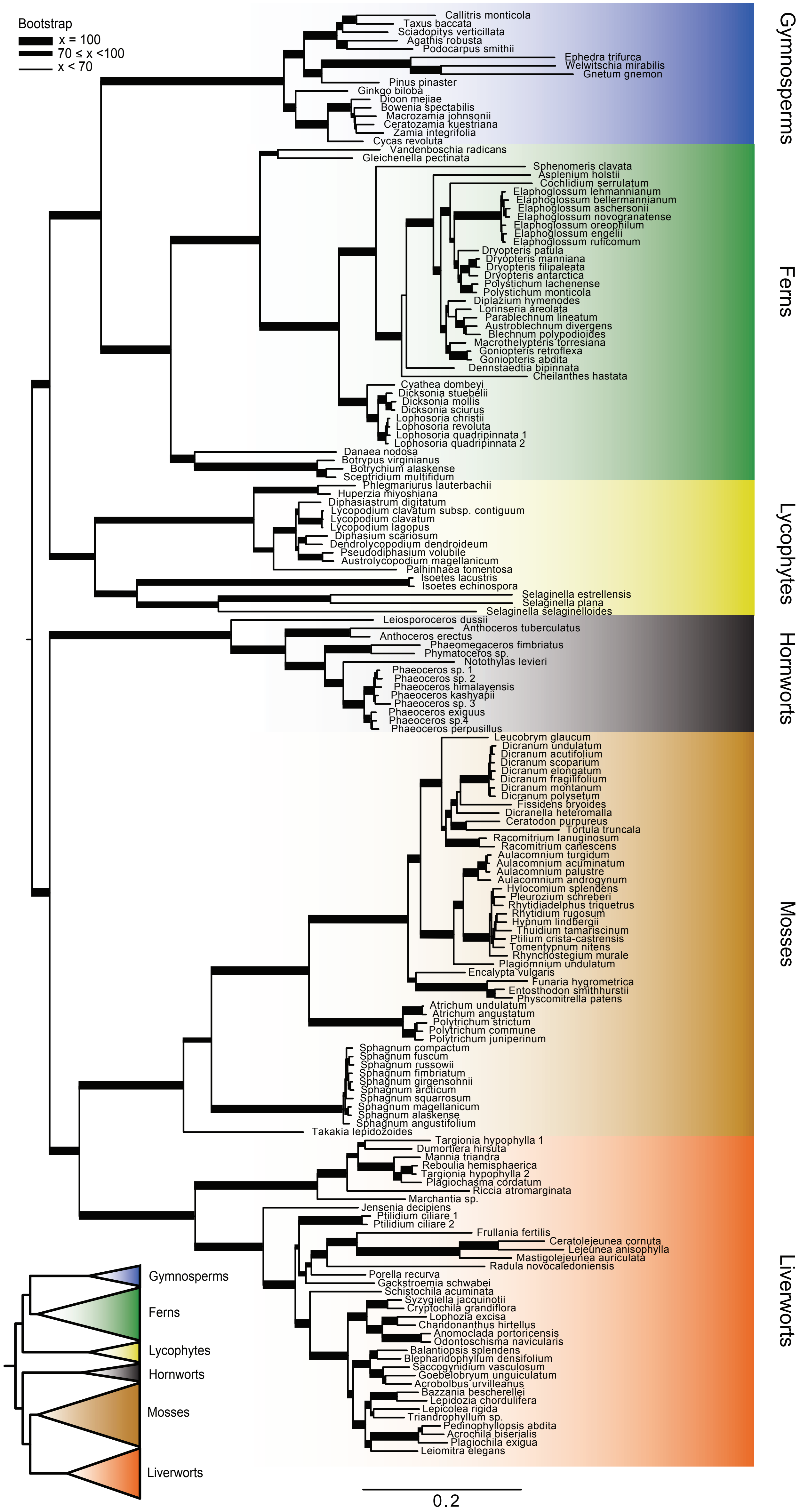
